# Modality specific roles for metabotropic GABAergic signaling and calcium induced calcium release mechanisms in regulating cold nociception

**DOI:** 10.1101/2022.05.12.491647

**Authors:** Atit A. Patel, Akira Sakurai, Nathaniel J. Himmel, Daniel N. Cox

**Affiliations:** Neuroscience Institute, Georgia State University, Atlanta, GA, USA

**Keywords:** CICR, GABA_B_, IP_3_R, RyR, nociception, thermosensation, sensory multimodality

## Abstract

Calcium (Ca^2+^) plays a pivotal role in modulating neuronal-mediated responses to multimodal sensory stimuli. Recent studies in *Drosophila* reveal class III (CIII) multidendritic (md) sensory neurons function as multimodal sensors regulating distinct behavioral responses to innocuous mechanical and nociceptive thermal stimuli. Functional analyses indicate that CIII-mediated multimodal behavioral output is dependent upon activation levels with stimulus-evoked Ca^2+^ displaying relatively low vs. high intracellular levels in response to gentle touch vs. noxious cold, respectively. However, the mechanistic bases underlying modality-specific differential Ca^2+^ responses in CIII neurons remain incompletely understood. We hypothesized that noxious cold-evoked high intracellular Ca^2+^ responses in CIII neurons may rely upon Ca^2+^-induced Ca^2+^ release (CICR) mechanisms involving transient receptor potential (TRP) channels and/or metabotropic G-protein coupled receptor (GPCR) activation to promote cold nociceptive behaviors. GABA_B_ receptor mutants and CIII-specific knockdown resulted in impaired noxious cold-evoked behaviors. Gαq and Phospholipase C signaling are likewise required for noxious cold sensing. Additionally, ER localized Ca^2+^ channels including the Ryanodine receptor (RyR) and Inositol trisphosphate receptor (IP_3_R) are required for cold nociceptive behaviors. GPCR mediated signaling, through GABA_B_-R2 and IP_3_R, is not required in CIII neurons for innocuous touch evoked behaviors. However, CICR via RyR is required in CIII neurons for innocuous touch-evoked behaviors. Disruptions in *GABA*_*B*_ *-R2, IP*_*3*_*R* and *RyR* in CIII neurons leads to significantly lower levels of cold-evoked Ca^2+^ responses indicating GPCR and CICR signaling mechanisms function in regulating Ca^2+^ release. CIII neurons exhibit bipartite cold-evoked firing patterns, where CIII neurons burst during rapid temperature change and tonically fire during steady state cold temperatures. *GABA*_*B*_*-R2* knockdown in CIII neurons resulted in disorganized firing patterns during cold exposure. Upon ryanodine pharmacological application, CIII neurons exhibit increased bursting activity and with CIII specific *RyR* knockdown, there is an increase in cold-evoked tonic firing and decrease in bursting. Lastly, our previous studies implicated the TRPP channel Pkd2 in cold nociception, and here, we show that *Pkd2* and *IP*_*3*_*R* genetically interact in regulating cold-evoked behavior. Collectively, these analyses support novel, modality-specific roles for metabotropic GABAergic signaling and CICR mechanisms in regulating intracellular Ca^2+^ levels and cold-evoked behavioral output from multimodal CIII neurons.

## 1 Introduction

Understanding how animals sense innocuous and/or potentially harmful stimuli, such as noxious temperature, chemical or mechanical insults, and respond appropriately is crucial for avoiding incipient damage that can lead to injury or death. Typically, upon sensing stimuli, an animal produces a set of behaviors that either mitigate or allow the animal to escape undesirable environmental conditions. Exposure to harmful temperatures can lead to impaired development, reproductive health, and survival, additionally, drastic changes in climate can lead to shifts in preferred temperature indices of species (Dowd et al., 2015; Williams et al., 2016). There are two types of metazoans: endotherms, which regulate body temperature independent of environment, and ectotherms, whose body temperature is dependent on environmental conditions (Clarke, 2014). Both endo- and ectotherms have adopted critical sensorimotor systems for maintaining thermal homeostasis and avoiding potentially harmful temperatures (Clarke, 2014; Gillooly et al., 2017). The somatosensory system plays a critical role in detecting thermal changes in vertebrates and invertebrates, where nociceptive neurons are strongly activated leading to tonic and/or bursting action potentials in response to potentially harmful stimuli (Himmel et al., 2017; Im & Galko, 2012). Integration and discrimination of sensory information is critical for animal survival, where failure to detect and avoid potentially harmful environmental cues can lead to injury. Understanding how integration and discrimination function in poly- or multi-modal sensory systems to generate stimulus relevant behavioral responses remain long-standing and active areas of research (Hu et al., 2017; Imambocus et al., 2022; Ohyama et al., 2015; Sherrington, 1906; Stockand & Eaton, 2013).

The *Drosophila melanogaster* larval peripheral nervous system primarily detects noxious (chemical, mechanical and/or thermal) stimuli through high threshold nociceptors innervating the barrier epidermis and are similar to mammalian unmyelinated c-fiber nociceptors. (Himmel et al., 2017; Im & Galko, 2012). There two primary nociceptive neuronal types, polymodal CIV md neurons, sensitive to noxious thermal (high heat), mechanical, and chemical stimuli, where neural activation leads to characteristic corkscrew body roll followed by rapid peristalsis in *Drosophila* larvae (Burgos et al., 2018; Himmel et al., 2019; Lopez-Bellido et al., 2019; Ohyama et al., 2013; Ohyama et al., 2015; Tracey et al., 2003; Vogelstein et al., 2014) and multimodal CIII md neurons that are sensitive to noxious thermal (cold) and innocuous mechanical stimuli (Jovanic et al., 2019; Kernan, 1994; Masson et al., 2020; Turner et al., 2016; Turner et al., 2018; Zhang et al., 2015). Detection of noxious (cold) and innocuous (mechanical) stimuli via CIII md neurons elicits stimulus specific behaviors.

Noxious cold evokes sustained contraction (CT) of head and tail, thus reducing overall surface area to volume ratio, while innocuous mechanical stimulation leads to a suite of behaviors including head withdrawal, pause, turning, and/or reverse locomotion (Jovanic et al., 2016; Kernan, 1994; Turner et al., 2016; Zhang et al., 2015). Sensory discrimination in CIII md neurons is dependent upon stimulus specific increases in cytosolic calcium (Ca^2+^) levels, where noxious cold stimulation leads to high levels of cytosolic Ca^2+^ and innocuous mechanical stimulation leads to mild increase of cytosolic Ca^2+^ (Turner et al., 2016). However, the mechanistic bases underlying modality-specific differential Ca^2+^ responses in CIII neurons remain incompletely understood.

In many animal species, transient receptor potential (TRP) channels have been implicated as primary sensory transducers for chemical, mechanical and thermal sensory modalities (Fowler & Montell, 2013; Himmel & Cox, 2020; Himmel et al., 2017; Liman et al., 2014; Patapoutian & Macpherson, 2006; Venkatachalam & Montell, 2007). Specifically, *Pkd2, Trpm* and *nompC* are required in cold sensitive CIII md neurons for noxious cold thermosensation and innocuous mechanosensation (Turner et al., 2016). Recent work has revealed chloride ion homeostasis and calcium-activated chloride channel function via the TMEM16/anoctamins, *subdued* and *white walker* (*wwk, CG15270*), as required for cold nociception but dispensable for detection of innocuous mechanical sensory information (Himmel, Sakurai, et al., 2021) providing initial clues to molecular bases of multimodal sensory discrimination in these neurons. Here, we sought to identify molecular bases of multimodal sensory discrimination that arise from differential Ca^2+^ dynamics in CIII md neurons. Therefore, we investigated signaling pathways involved in increasing cytosolic Ca^2+^ concentrations. Intracellular increases in cytosolic Ca^2+^ can result from both ionotropic and metabotropic pathways. Ionotropic pathways include extracellular Ca^2+^ influx from voltage-gated Ca^2+^ channels and TRP channels localized on the plasma membrane and organellar Ca^2+^ efflux from mitochondria and endoplasmic reticulum (ER) (Berridge, 1998; Fowler & Montell, 2013; Grienberger & Konnerth, 2012; Turner et al., 2016; Verkhratsky & Petersen, 1998). Metabotropic pathways, specifically G protein coupled receptors (GPCRs), are involved in a variety of sensory systems including vision, olfaction, audition, and thermosensation, and in part, function through organellar Ca^2+^ release (Fukuto et al., 2004; Gu et al., 2022; Herman et al., 2018; Im et al., 2015; Imambocus et al., 2022; Jin et al., 2013; Julius & Nathans, 2012; Leung & Montell, 2017). ER luminal Ca^2+^ efflux into the cytosol functions through the Ca^2+^ induced Ca^2+^ release (CICR) pathway via ryanodine receptor (RyR) and inositol trisphosphate receptor (IP_3_R). We hypothesize that noxious cold-evoked high intracellular Ca^2+^ responses in CIII neurons may rely upon CICR mechanisms via transient receptor potential (TRP) channels and/or metabotropic GPCR activation to promote cold nociceptive behaviors.

In this study, we found that metabotropic GPCR and CICR signaling pathway genes are expressed in CIII md neurons and are required for cold nociceptive behaviors in third instar *Drosophila* larvae. Impaired GABA_B_-R2 receptor, RyR and IP_3_R signaling leads to reductions in cold-evoked Ca^2+^ responses. Extracellular electrical recordings of CIII md neurons revealed GABA_B_-R2 is required for proper patterning of cold-evoked activity and cold-evoked RyR mediated Ca^2+^ dynamics promote neuronal bursting. GABA_B_-R2 receptor and IP_3_R signaling is required specifically for cold nociception but not innocuous mechanical sensation, whereas RyR is required for both innocuous mechanical sensation and noxious cold detection. GABA_B_-R2 receptor, RyR and IP_3_R signaling are not, however, required for CIII md neuron dendrite morphogenesis, action potential propagation or neurotransmitter release indicative of roles for these signaling machinery at the sensory transduction stage.

## 2 Methods

### 2.1 Fly Strains

All *Drosophila melanogaster* strains used in this study are listed in (**Supplementary Table 1**). All *Drosophila* reagents were maintained on standard cornmeal diet in 12:12 hour light-dark cycle at ~22°C. All experimental crosses were raised in 12:12 hour light-dark cycle at ~29°C, unless otherwise stated. We utilized several class III neuron *GAL4* drivers: *19-12*^*GAL4*^ (Xiang et al., 2010), *nompC*^*GAL4*^, and *GMR83B04*^*GAL4*^.

### 2.2 Cold Plate Assay

We assessed *Drosophila melanogaster* larval responses to noxious cold temperatures via cold plate assay (Turner et al., 2016; Patel and Cox, 2017), where we acutely expose third instar larvae to cold temperatures (**Figure 2-2**).

#### Equipment

We used Nikon DSLR (D5300) for recording larval behavioral response. The camera was mounted on a tripod, where the DSLR was facing down towards the table and focused on a pre-chilled (10°C) TE technologies Peltier plate (CP-031, TC-48-20, RS-100-12). We used goose neck oblique light source (Nikon, NI-150) to increase signal to noise ratio for better data processing.

#### Experiment

All F1 genetic crosses progeny used in assessing cold-evoked responses were raised in 29°C incubator. We tested age matched third instar larvae. We gently removed larvae from food using a thin brush and placed them on a wet Kimwipe. Next, food debris was passively removed by allowing the animals to freely locomote and then gently placing them on secondary wet Kimwipe. Using a spray bottle, we lightly misted water onto a thin black metal plate, after which we placed 5-7 larvae onto the plate. We allowed larvae to resume locomotion, while removing any excess water droplets from around the animals. Next, we began video recording on DSLR (1920×1080 pixels at 30 frames per seconds) and we placed the thin metal plate on a pre-chilled Peltier device. We assessed cold-evoked behaviors for first five seconds of stimulus.

Semi-automatic video processing and behavioral quantification: For high throughput video processing, we used FIJI and Noldus Ethovision XT. We used video to video convertor (https://www.videotovideo.org/) for uncompressing raw video files. We manually selected the first frame at which cold stimulus was delivered. Next, using custom built FIJI macro we processed the videos. The custom macro first trims each video to desired duration, identifies individual larvae using automatic thresholding (Triangle method) and creates individually cropped (250×250pixels) larval videos. For increased throughput and signal to noise ratio, we next automatically removed background from individually cropped larval videos using automatic thresholding (Li method). For semi-automated quantitative surface area measurements, we utilized Noldus Ethovision XT software, only larvae that were elongated at the onset of the first frame were analyzed. For automatic data processing, we used custom built R macros. Briefly, custom R macro combined raw surface area measurements for individual animals within each genotype. Next, we calculated percent change in surface area over time, duration of response, and magnitude of behavioral response. Our primary threshold for behavioral analysis, derived from an in-depth examination of the behavior, percent contraction entails a response of at least a 10% reduction in surface area from initial area for at least 0.5 seconds. From this metric we also derived CT duration, the total time an animal spends below −10% change in surface area, and CT magnitude, defined as average percent change in surface area for duration of the assay.

#### Statistical analysis

We preformed following statistical tests for all cold plate assay data analysis. %CT response: Fisher’s exact with Benjamini-Hochberg for multiple comparison. We used r for performing comparisons of percent behavior response between genotypes, we used Benjamini-Hochberg multiple comparison correction. CT duration: Kruskal-Wallis with Benjamini, Krieger and Yekutieli for multiple comparisons. CT duration data are not normal. CT magnitude: One-way ANOVA with Holm-Šídák’s for multiple comparisons.

### 2.3 Confocal Imaging

*In vivo* dendrite morphology imaging: For assessing md CIII dendrite morphology, we crossed class III specific driver driving membrane tagged GFP (*nompC*^*Gal4*^*> mCD4::tdGFP*) to gene specific UAS-RNAi lines. Aged matched live third instar larvae were mounted on a microscope slide and anesthetized by using few drops of halocarbon and ether solution (1:5 v/v). Three-dimensional z-stacks were imaged using Zeiss LSM780 with Plan-Apochromat 20x/0.8 M27 objective and 488nm laser. Image dimensions were 1024 by 1024 pixels (607.28µm x 607.28µm) with 2µm step size.

Quantitative dendrite morphology analysis was performed as previously described in (Das et al., 2017). GABA_B_-R2 expression imaging: For visualizing GABA_B_-R2 expression in md neurons, we utilized a recently developed *Drosophila* reagent for CRISPR mediated insertion of RFP after the first exon in *GABA*_*B*_*-R2* (Bloomington Stock# 84504). We visualized expression of *GABA*_*B*_*-R2::RF*P in md CIII neurons using *nompC*^*Gal4*^*> mCD4::tdGFP*. We imaged the third instar larvae similar to dendrite morphology imaging. We captured red fluorescence using 561nm laser. We identified md CIII neurons using GFP signal, and we identified other md and es neurons based on cell body position. Red and green fluorescence intensity were analyzed using Zeiss Zen blue software.

### 2.4 Calcium imaging assays

#### GCaMP imaging

We utilized GCaMP6m for analyzing transient *in vivo* cold-evoked Ca^2+^ dynamics in class III md neurons. We mounted live intact third instar larvae expressing GCaMP6m and RNAi in CIII md neurons (*19-12*^*GAL4*^) onto microscope slide as described previously (Patel & Cox, 2017; Turner et al., 2016). We placed the microscope slide onto a small peltier stage (Linkam Scientific Instruments, Linkam Scientific, model: PE120, T95 linkpad and PE95) mounted on the microscope. Using Zeiss LSM780 laser confocal microscope, we imaged CIII md neuron (ddaF) in larval abdominal segment 1. After mounting the animal and visualizing the neuron of interest, we allowed the larva to acclimate to the set up and allowed calcium levels to return baseline. To mitigate condensation on the coverslip created from cold exposure, a very light air current was directed on top of the coverslip and below the objective lens. We next acquired time-lapse data (~13frames per second at 62.51µm x 62.51µm resolution) and with the following stimulus paradigm (30 seconds baseline (25°C), ramp down to 6°C at 20°C per minute, 10 seconds at 6°C, ramp up baseline at 20°C per minute, and hold at baseline for 30 seconds).

#### Quantitative GCaMP analysis

We stabilized raw time-lapse videos using Rigid Body transformation from StackReg in FIJI. (Linkert et al., 2010; Schindelin et al., 2012; Thevenaz et al., 1998) We manually drew region of interest (ROI) around the cell body and measured area normalized fluorescence for each frame. Next, we calculated changes in fluorescence from baseline using the following equation: 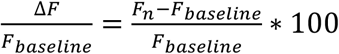 We report normalized ΔF/F_baseline_ overtime and report average peak ΔF/F_baseline_.

#### CaMPARI assay and imaging

We utilized a Ca^2+^ integrator CaMPARI2 for assessing cold-evoked Ca^2+^ levels at a larger neural population level as previously described (Moeyaert et al., 2018; Patel & Cox, 2017; Turner et al., 2016). We expressed *CaMPARI2* in CIII md neurons (*GMR83B04*^*GAL4*^) along with RNAi and tested age matched third instar larvae. We used our cold plate assay Peltier plate for delivering noxious cold stimulus and used AxioScope V16 Zeiss microscope for delivering photoconverting light. We placed individual third instar larvae on lightly water misted black metal plate. We allowed the animals to acclimate to plate before placing the black metal plate on the pre-chilled (experimental) or room temperature (control) Peltier plate, while simultaneously exposing the animals to photoconverting light. We exposed the larvae to cold and photoconverting light for 20 seconds. Next, within five minutes, we mounted the animals on a microscope slide and imaged dorsal clusters of CIII md neurons in abdominal segments 1-3. Imaging was conducted similarly as described in dendrite morphology imaging section.

#### CaMPARI quantitative analysis

For analyzing CaMPARI2 data, we chose a dual approach to analyze the red to green ratio from CaMPARI2 signal at the cell bodies and perform a modified Sholl analysis, where we measure fluorescence intensity at radial distances from the cell body.

#### Cell body analysis

We created a custom automatic pipeline for drawing regions of interest, where users verify ROIs before fluorescence quantification. Using FIJI, we created a custom script, where a maximum intensity projection of each z-stack image is created. Next, the GFP signal was thresholded (Moments method), then background and branches are removed using erode and dilate functions. After background clearing only the cell bodies remain, and the Analyze particle function is used to draw ROIs around cell bodies. We next created a separate batch processing macro, where we manually verify the accuracy of each ROI and manually redraw any incorrect ROIs. Upon ROI verification, area normalized red and green fluorescence intensities were extracted. As previously described (Fosque et al., 2015; Im et al., 2018; Patel & Cox, 2017; Turner et al., 2016), we report evoked photoconverted CaMPARI signal as F_green_/F_red_ ratio.

#### Sholl intensity analysis

We created a set of sequential macros using FIJI for clearing background, drawing radial ROIs for cell body and dendrites of interest, and measuring fluorescence intensities at each radius. Briefly, for background clearing we manually threshold the GFP signal, remove neurites from neighboring cells using the “Paintbrush tool”, and using the “Wand tool” to select all branches and cell body for the neuron of interest. In a batch processing macro, which draws five-pixel wide radial ROIs at a single pixel interval, where only the dendrites and cell body are selected. After radial ROIs are drawn, area and area normalized red and green fluorescence intensities are quantified for each radial step from the center of the cell body. We report evoked photoconverted CaMPARI2 signal as F_green_/F_red_ ratios away from the center of the cell body.

### 2.5 Optogenetic Assay

We tested neural excitability and neural transmission capabilities for class III md neurons with *RyR, IP*_*3*_*R* or *GABA*_*B*_*-R2* knockdown. We crossed each UAS-RNAi line with class III specific optogenetic line (*19-12*^*GAL4*^*>ChETA::YFP*). *Drosophila* do not produce a requisite light sensitive cofactor, all trans-retinal (ATR). Therefore, all adult animals in a cross were placed in ATR supplemented food, where the final ATR concentration in food plug was 1500µM. All F1 progeny were raised in ATR supplement food. All crosses were kept in a dark box at 29°C.

#### Equipment

We created a custom rig for our optogenetic experiments, where we implemented principles of dark field illumination for high signal to noise ratio imaging of *Drosophila* larval behaviors. We used a ring stand to mount white led ring light, where we limited white light exposure to on 2.5cm circular opening on the stage. We placed two Thorlabs 470nm (M470L3-C4) led lights below the stage. We recorded animals from above using Canon T3i DSLR camera. We used Noldus Ethovision XT software and Thorlabs led systems for automatically recording and controlling blue light exposure. (Thorlabs: DC4100, DC4100-hub, and two M470L3-C4 led light. Noldus: mini-IO box)

#### Experimental approach

We tested age matched third instar larvae for blue light evoked responses in a dark room. Using a brush, we gently picked out third instar larvae from food and placed them on wet Kimwipe. Next, the animals were placed on a lightly water misted thin clear glass plate. The glass plate was placed onto the optogenetic ring stage and animals were exposed to blue light. Our neural stimulation paradigm was five seconds baseline (blue light off), 10 seconds of neural activation (blue light on) and five seconds neural inactivation (blue light off).

#### Quantification and analysis

*Drosophila* larval surface area was measured live using Noldus Ethovision XT. We used similar data processing pipeline as for cold plate assay analysis including percent response, duration, and percent change in area over time.

### 2.6 Microarray analysis

We analyzed our previously published microarray datasets, which were deposited in Gene Expression Omnibus, md CIII (GSE69353), md CIV (GSE46154), and whole larval (GSE46154) microarray datasets from Cox lab (Iyer et al., 2013; Turner et al., 2016).

### 2.7 Innocuous mechanical assay

We tested age matched third instar *Drosophila* larvae for behavioral responses to innocuous mechanical stimulation. (Himmel, Sakurai, et al., 2021; Kernan, 1994; Turner et al., 2016) We placed third instar larvae on lightly misted black metal plate, same as cold plate assay, and allowed the animals to acclimate to the new arena. Using a single brush bristle, we gently touch the anterior segments. We performed three separate simulations with 30 seconds between each trial. We observed animal responses via Zeiss Stemi 305 microscope. After each mechanical stimulation, we manually scored the following behaviors: no response, pausing, head withdrawal, turning, and reverse locomotion. Each behavioral response receives a score of 1 and no response receives 0. We totaled scores from three trials for a max total of 12 and report average scores for each genotype.

Additionally, we analyzed single trial total score (max 4), where we report proportion of each score in each trial for individual genotypes.

### 2.8 Electrophysiology

Extracellular recordings of age-matched third instar larvae were performed in de-muscled filleted preparations submerged in physiological saline and exposed to cold temperatures.

#### Larval preparation

A third instar *Drosophila* larva was dissected using fine scissors along the ventral midline and was fixed to the bottom of the experimental chamber by pinning at the edge of the body wall of the fillet. After gently removing internal organs, longitudinal muscles were all removed by using a tungsten needle and fine scissors. The experimental chamber was filled with physiological saline (200µL) and perfused at 30-40µL/sec. The composition of saline is as follows in mM: NaCl 120, KCl 3, MgCl2 4, CaCl2 1.5, NaHCO3 10, trehalose 10, glucose 10, TES 5, sucrose 10, HEPES 10. (Xiang et al., 2010)

#### Temperature control

Cold temperature stimulus was delivered using an in-line solution cooler (SC-20, Warner Instruments, Hamden, CT, USA) and controlled by single channel temperature controller (CL-100, Warner Instruments, Hamden, CT, USA). The temperature in experimental chamber was constantly monitored by placing microprobe thermometer (BAT-12, Physitemp, Clifton, NJ, USA) close to larval fillet.

#### Extracellular recording

Class III md neuronal extracellular activity was recorded using a borosilicate glass micropipette electrode (tip diameter, 5-10µM) by applying gentle suction. Voltage clamp recordings were obtained using patch-clamp amplifier (MultiClamp 700A, Molecular Devices, San Jose, CA, USA). The patch-clamp amplifier output was digitized at 10kHz sampling rate using a Micro1401A/D (Cambridge Electronic Design, Milton, Cambridge, UK) and imported into a laptop using Spike2 software version 8 (Cambridge Electronic Design, Milton, Cambridge, UK).

For gene knockdown experiments, we expressed *GABA*_*B*_*-R2*^*RNAi*^ or *RyR*^*RNAi*^ in GFP tagged class III md neurons using *19-12*^*GAL4*^. Extracellular recordings of ddaF md CIII neurons were obtained during cold exposure. The following temperature paradigm was used: 30 seconds at room temperature (~22°C), fast ramp down to stimulus temperature (−3.3 ± 0.86 °C/s, mean ± SD) and hold for 60 seconds, and fast ramp up to room temperature (−3.3 ± 0.86 °C/s, mean ± SD). We tested three stimulus temperatures (20°C, 15°C and 10°C), where stimulus delivery was randomized between animals and there was >2 min interval between trials.

For ryanodine pharmacology, we used GFP tagged CIII ddaF neurons (*19-12>GFP*) and applied cold exposure similar to gene knockdown electrophysiological recordings. Ryanodine (10^−6^ M) was applied through the perfusion path.

Spike sorting and firing rate classification: After each experiment, spikes were detected off-line by setting a threshold current in Spike2 software to calculate the average spiking rate in each time bin (10 sec). For bursting spikes, a group of three or more consecutive spikes with a spike interval of less than 0.15 seconds was considered as a burst, and the spikes that constitute each burst were defined as bursting spikes. The remaining spikes were defined as tonic spikes.

### 2.9 Statistical analysis and data visualization

Statistical analyses were performed using R (Fisher’s exact test) and GraphPad Prism. All data were graphed using GraphPad Prism.

## 3 Results

### 3.1 Metabotropic GABA receptors are required for cold nociception

CIII md neurons differentially encode mechanical and thermal stimuli by utilizing differential activation levels mediated by Ca^2+^ signaling (Turner et al., 2016). Innocuous mechanical stimulation results in low evoked cytosolic Ca^2+^ responses, whereas noxious cold temperature results in high stimulus evoked cytosolic Ca^2+^ responses. Previous work has implicated the transient receptor potential (TRP) channels Pkd2, NompC, and Trpm in sensing innocuous mechanical stimuli and noxious cold (Turner et al., 2016), however, the molecular bases of how CIII md neurons discriminately encode modality-specific information to elicit stimulus-relevant behavioral output remain unclear. We hypothesize that differential cytosolic Ca^2+^ levels in CIII md neurons likely arise from signaling mechanisms involved in cytosolic Ca^2+^ dynamics. Therefore, we first investigated the roles of G protein coupled receptors (GPCR) in noxious cold-evoked *Drosophila* larval behaviors.

We leveraged our previous cell type specific transcriptomic datasets to identify GPCRs that might be required in cold sensing specifically in CIII md neurons. We analyzed ionotropic GABA_A_ (Rdl) and metabotropic GABA_B_ (-R1, -R2, -R3) receptor expression levels in CIII md neurons, where on average Rdl, GABA_B_-R1, and GABA_B_-R3 had lower expression compared GABA_B_-R2 (**Figure 1A**). Additionally, we analyzed metabotropic GABA_B_ (-R1, -R2, -R3) receptor expression for enrichment in CIII neurons by comparing expression levels between CIII md neurons, CIV md neurons, and whole larvae (WL). We have previously demonstrated that CIV neurons are largely cold-insensitive and are not required for cold-evoked CT behavior (Turner et al., 2016). CIII md neuron GABA_B_-R1 receptor expression was 1.8-fold higher compared to CIV md neurons and 0.26-fold lower compared to WL (**Figure 1B**). CIII md neuron expression for GABA_B_-R2 and -R3 receptor were similar, with an ~4-fold higher expression compared to CIV md neurons and ~2-fold higher expression compared to WL (**Figure 1B**). To further validate our GABA_B_-R2 CIII md neuron expression data, we visualized GABA_B_-R2 receptor localization. In order to visualize GABA_B_-R2 expression, we took advantage of recently published CRISPR-mediated fluorescently tagged GABA_B_-R2 receptor, where red fluorescent protein (RFP) was inserted after the first exon (Deng et al., 2019). We co-expressed membrane tagged green fluorescent protein (mCD8::GFP) and GABA_B_-R2::RFP in CIII md neuron using the *nompC*^*GAL4*^ driver. Live confocal imaging data revealed that GABA_B_-R2::RFP expression is not limited to CIII md neurons, but extended to other PNS neurons including multiple md neuron subtypes (CI & CII) and external sensory (ES) neurons (**Figure 1C**). Interestingly, CII md neurons have the highest GABA_B_-R2::RFP expression, followed by ES and CI md neurons with CIII md neurons showing the lowest relative expression (**Figure 1C**). Collectively, transcriptome data reveal that GABA_B_-R2 receptor has the highest expression levels of the three metabotropic GABA receptors and GABA_B_-R2::RFP receptor expression is present on dendrites and the cell body of CIII md neurons.

**Figure 1:**
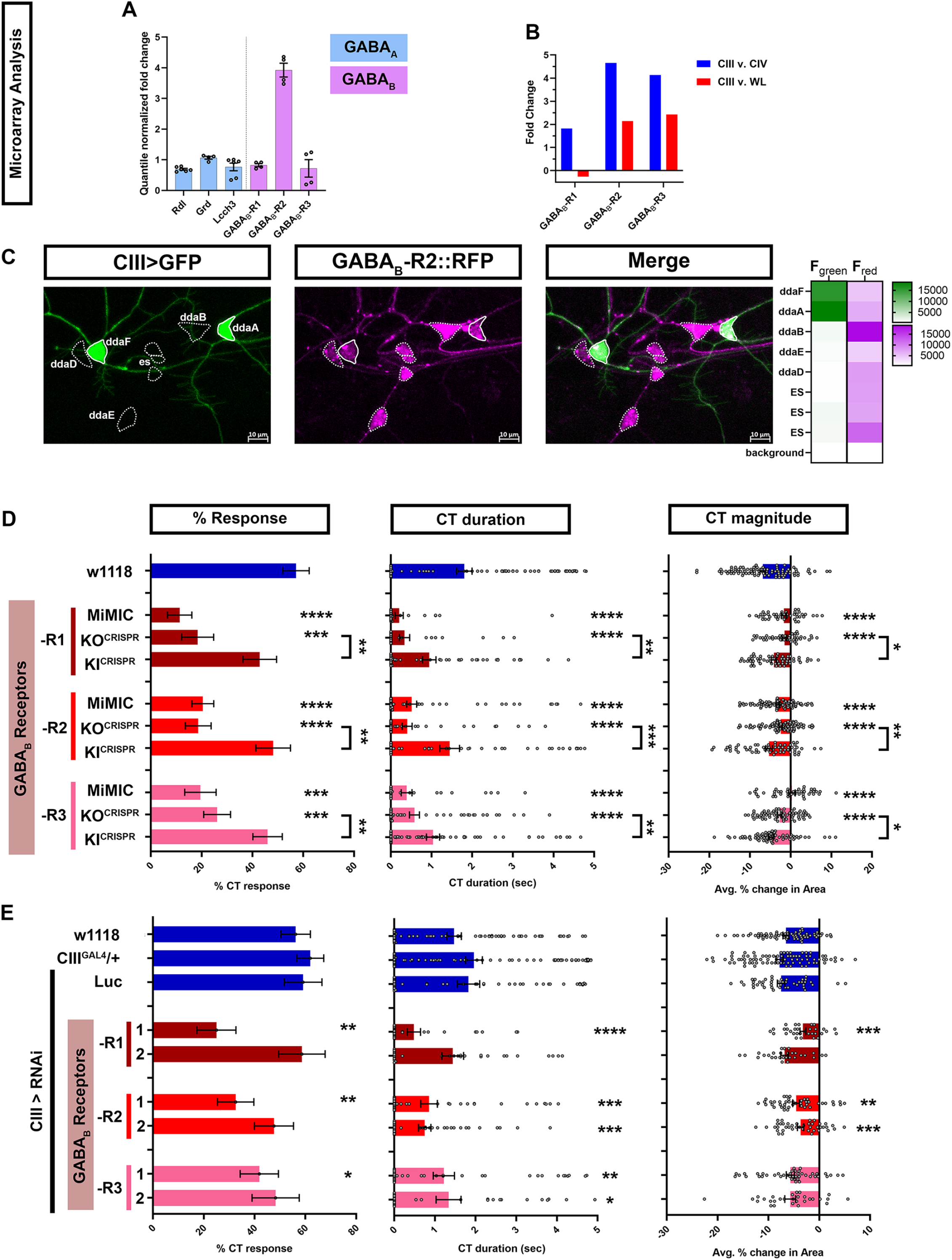
Metabotropic GABA_B_ are expressed in CIII md neurons and are required for cold-evoked CT behavior. (**A-B**) md neuron transcriptomic analysis. (**A**) Quantile normalized expression levels of ionotropic (*GABA*_*A*_) and metabotropic (*GABA*_*B*_) receptors in CIII md neurons. All receptors are expressed above background levels and *GABA*_*B*_*-R2* has very high expression in CIII md neurons. (**B**) Comparisons of transcriptomes between CIII md neurons and CIV md neurons or whole larvae (WL). We focused on metabotropic GABA receptors, which show, in general, greater expression in CIII md neurons compared to CIV md neurons or whole larvae (WL). (**C**) Visualization of GABA_B_-R2 expression in *Drosophila* larval PNS. CIII md neurons were visualized using membrane anchored GFP expression to mark overall morphology and GABA_B_-R2 expression was visualized using an RFP tagged GABA_B_-R2. GABA_B_-R2 RFP expression is detectable in CIII (ddaF, ddaA), CII (ddaB), CI (ddaD, ddaE) and ES neurons. Genotype: GABA_B_-R2::RFP, *nompC*^*GAL4*^>*GFP*. (**D-E**) Cold plate assay assessing cold-evoked behaviors of third instar *Drosophila* larvae. We report %CT (left), CT duration in seconds (middle), CT magnitude (right). (**D**) Whole animal mutant analysis of GABA_B_ receptor’s role in cold-evoked behaviors. For each GABA_B_ (-R1, -R2, -R3) receptor, we tested one MiMIC and one CRISPR knockout (KO) line (truncated after first exon and inserted RFP). Additionally, we tested CRISPR GAL4 knock-in (KI) line, which replaces the RFP of KO^CRISPR^ with the remaining exons restoring receptor function. N_Average_ = 65. (**E**) Class III specific knockdown of *GABA*_*B*_ *(-R1, -R2, -R3)* receptors. Control conditions include: *w1118, 19-12*^*GAL4*^*/+*, and *19-12>Luc*^*RNAi*^. There is no statistical difference between the controls. Knockdown of *GABA*_*B*_ (*-R1, -R2, -R3*) receptors was tested with two independent RNAi lines and comparisons made to *19-12*^*GAL4*^*/+* control. N_Average_ = 46. Significant differences indicated via asterisks, where *p<0.05, **p<0.01, ***p<0.001, and ****p<0.0001.

To assess the role of metabotropic GABA_B_ receptors, we exposed the ventral surface of *Drosophila* larvae to acute noxious cold temperatures (≤10°C) using a cold plate assay and observed stimulus evoked behavioral responses. *Drosophila* larvae have a stereotyped bilateral (anterior & posterior) withdrawal towards the middle of the animal, we defined this behavior as contraction (CT) response (**Supplementary Figure 1**) (Patel & Cox, 2017; Turner et al., 2016). Cold plate assays performed on mutant larvae carrying Minos-mediated integration cassette (MiMIC) gene traps for GABA_B_ receptors revealed significant deficits in cold-evoked CT response (**Figure 1D**). Only heterozygotes were viable for *GABA*_*B*_*-R1* and *GABA*_*B*_*-R2* MiMICs, which had 11% and 20% CT response, respectively, compared to 57% CT response for genetic controls (*w1118*). Homozygous *GABA*_*B*_*-R3* MiMIC mutant larvae also had significant reduction in %CT response compared to control (19% v. 57% CT response). *GABA*_*B*_ (*-R1, -R2, -R3*) MiMICs all exhibited significant reductions in CT duration and magnitude of CT response compared to controls (**Figure 1D**). To independently validate our results for *GABA*_*B*_ MiMICs, we used recently published GABA_B_ truncation mutants (referred to as *GABA*_*B*_*-KO*) fused to RFP (Deng et al., 2019). *GABA*_*B*_ (*-R1, -R2, -R3*) KO mutants all displayed significant reductions in CT response, CT duration and CT magnitude compared to control (**Figure 1D**), similar to what we observed for *GABA*_*B*_ (*-R1, -R2, -R3*) MiMICs. We also used corresponding genomic rescue constructs (referred to as *GABA*_*B*_*-KI*), where truncation mutants (*GABA*_*B*_*-KO*) were used to generate knock-ins by reinserting truncated exons and fusing GAL4 to the C-terminus and tested whether *GABA*_*B*_*-KI* would rescue noxious cold sensing deficits observed in GABA_B_-KO mutants. On their own, all *GABA*_*B*_ (*-R1, -R2, -R3*) KI lines had mild, but insignificant reductions in cold-evoked CT behavioral response relative to control (**Figure 1D**). However, all *GABA*_*B*_ (*-R1, -R2, -R3*) KI lines were sufficient to significantly rescue *GABA*_*B*_ (*-R1, -R2, -R3*) KO mutant phenotypes for all the parameters tested (**Figure 1D**). Collectively, our whole animal mutant data suggest that GABA_B_ (-R1, -R2, -R3) are required for noxious cold sensing in *Drosophila* larvae. We next drove expression of *GABA*_*B*_^*RNAi*^ using a cell type specific *GAL4* driver for CIII md neurons (*19-12*^*GAL4*^) and assessed cold-evoked behavioral responses. We controlled for genetic background (*w1118*), GAL4 expression (*CIII*^*GAL4*^/+), and RNAi activity (*LUC*^*RNAi*^). All three control conditions had relatively similar %CT response, CT duration and CT magnitude. Only one of the two independent gene-specific RNAi lines tested for *GABA*_*B*_*-R1, -R2*, or *-R3* gene knockdown led to significant reductions in %CT response compared to controls (**Figure 1E**). *GABA*_*B*_*-R1* knockdown in CIII md neurons showed significant reductions in CT duration and magnitude in only one of the two RNAi lines tested. However, both RNAi lines tested for *GABA*_*B*_*-R2* knockdown in CIII md neurons led to significant reductions in CT magnitude and duration (**Figure 1E**). *GABA*_*B*_*-R3* knockdown in CIII md neurons had the mildest effect on cold-evoked CT behavioral response with only significant reductions in CT duration for both RNAi lines tested (**Figure 1E**). Whole animal mutations in GABA_B_ receptors showed decreased cold sensitivity and cold sensitivity was rescued for all three GABA_B_ receptors using KI^CRISPR^ constructs. Consistent with mutant analyses, our cell-type specific knockdowns of GABA_B_-R1 or -R2 led to strong reductions in cold-evoked responses, however, only GABA_B_-R2 showed similar deficits in cold-evoked responses using two independent RNAi lines. GABA_B_-R1 and -R2 form heterodimers, where GABA_B_-R1 contains GABA binding site and GABA_B_-R2 associates with heterotrimeric G-proteins (Benarroch, 2012; Papasergi-Scott et al., 2020). Collectively, GABA_B_-R2 is highly enriched in CIII md neurons compared to other GABA receptors, impairments in GABA_B_-R2 signaling showed consistent phenotypes for both whole animal mutants and cell-specific gene knockdown in CIII md neurons, and GABA_B_-R2 binds to heterotrimeric G-proteins, therefore, we primarily focus on GABA_B_-R2 for the remainder of the analyses.

### 3.2 Calcium induced calcium release mechanisms are necessary for cold sensing

We next sought to investigate signaling mechanisms involved in stimulus specific neural activation levels of CIII md neurons through increases in cytosolic Ca^2+^. Noxious cold stimulus significantly increases cytosolic Ca^2+^ levels through Ca^2+^ permeant TRP channels which are required for proper cold-evoked CT responses (Turner et al., 2016). Apart from Ca^2+^ influx through TRP channels, another possible mechanism for elevating cytosolic Ca^2+^ is from intracellular stores mediated through GPCR downstream signaling.

Activated GPCR’s may function through variety of trimeric G proteins including G_αi/o_, G_αq_, and G_αs_. G_αi/o_ and G_αs_ generally function in opposing directions, whereby G_αi/o_ inhibits adenylyl cyclase to reduce cAMP and G_αs_ activates adenylyl cyclase to increase cAMP. G_αq_ activation leads to phospholipase Cβ (PLC) hydrolyzing phosphatidylinositol 4,5-bisphosphate (PIP_2_) to diacylglycerol (DAG) and inositol 1,4,5-trisphosphate (IP_3_). Furthermore, IP_3_ production leads to activation of IP3 receptors (IP_3_R) releasing endoplasmic reticulum (ER) luminal Ca^2+^ thereby increasing cytosolic Ca^2+^ levels. In rat hippocampal neuron cultures, GABA_B_ mediated increases in Ca^2+^ current is dependent upon G_αq_ signaling, however, it is unknown whether PLC activation is required (Karls & Mynlieff, 2015). Additionally, GABA_B_ receptors have previously been shown to co-localize with G_αq_ (Karls & Mynlieff, 2015). Therefore, we investigated whether the G_αq_ signaling pathway is required for *Drosophila* larval cold nociception.

The *Drosophila melanogaster* genome encodes only one *G*_*αq*_ gene and three PLC genes (*norpA, small wing* (*sl*), and *Plc21C*). Our CIII md neuron transcriptome analysis revealed that *G*_*αq*_, *norpA, Plc21C* and *sl* are expressed more than 1.25-fold compared to either CIV md neurons or whole larva (**Supplementary Figure 2A**). Next, we assessed the requirement for G_αq_ and PLCs in CIII md neurons for cold nociception. Two independent CIII md neuron specific knockdowns of G_αq_ led to significant decreases in %CT response, CT duration and magnitude compared to controls. In addition, *norpA, sl*, and *Plc21C* gene knockdowns in CIII md neurons led to significant reductions in cold-evoked %CT response, CT duration and magnitude compared to controls (**Supplementary Figure 2B**). In summary, G_αq_ and PLCs are required for cold-evoked behaviors.

TRP channels have been implicated in cold-evoked increases in cytosolic Ca^2+^ and we have shown evidence for metabotropic GABA signaling being required for cold nociception (Turner et al., 2016). Cytosolic Ca^2+^ concentrations are tightly controlled, where baseline concentrations range from 0.1-1µM. By contrast, baseline ER luminal Ca^2+^ concentrations are maintained at much higher concentrations (10^−3^M) (Reddish et al., 2021). Plasma and ER membranes share close apposition to one another in neurons at cell body, dendrites, and axons, where Ca^2+^ second messenger signaling can be tightly regulated (Woll & Van Petegem, 2022; Wu et al., 2017). IP_3_R and RyR are two major Ca^2+^ ion channels located on ER membrane, where both receptors are activated by Ca^2+^ binding and additionally IP_3_R is also gated by IP_3_ (Woll & Van Petegem, 2022). Therefore, we hypothesize stimulus evoked TRP channel mediated increases in Ca^2+^ result in activation of CICR mechanism and/or GPCR signaling results in Ca^2+^ release from ER stores.

Class III md neuron transcriptomics reveals relatively high levels of RyR and IP_3_R expression above background (**Figure 2A**). Comparative md neuron transcriptome analyses revealed that both IP_3_R and RyR are highly enriched in CIII md neurons compared to CIV md neurons (**Figure 2B**). RyR expression in CIII md neurons is 42-fold higher than whole larvae and IP_3_R expression in CIII md neurons is modestly enriched at 3.5-fold compared to whole larva (**Figure 2B**).

**Figure 2:**
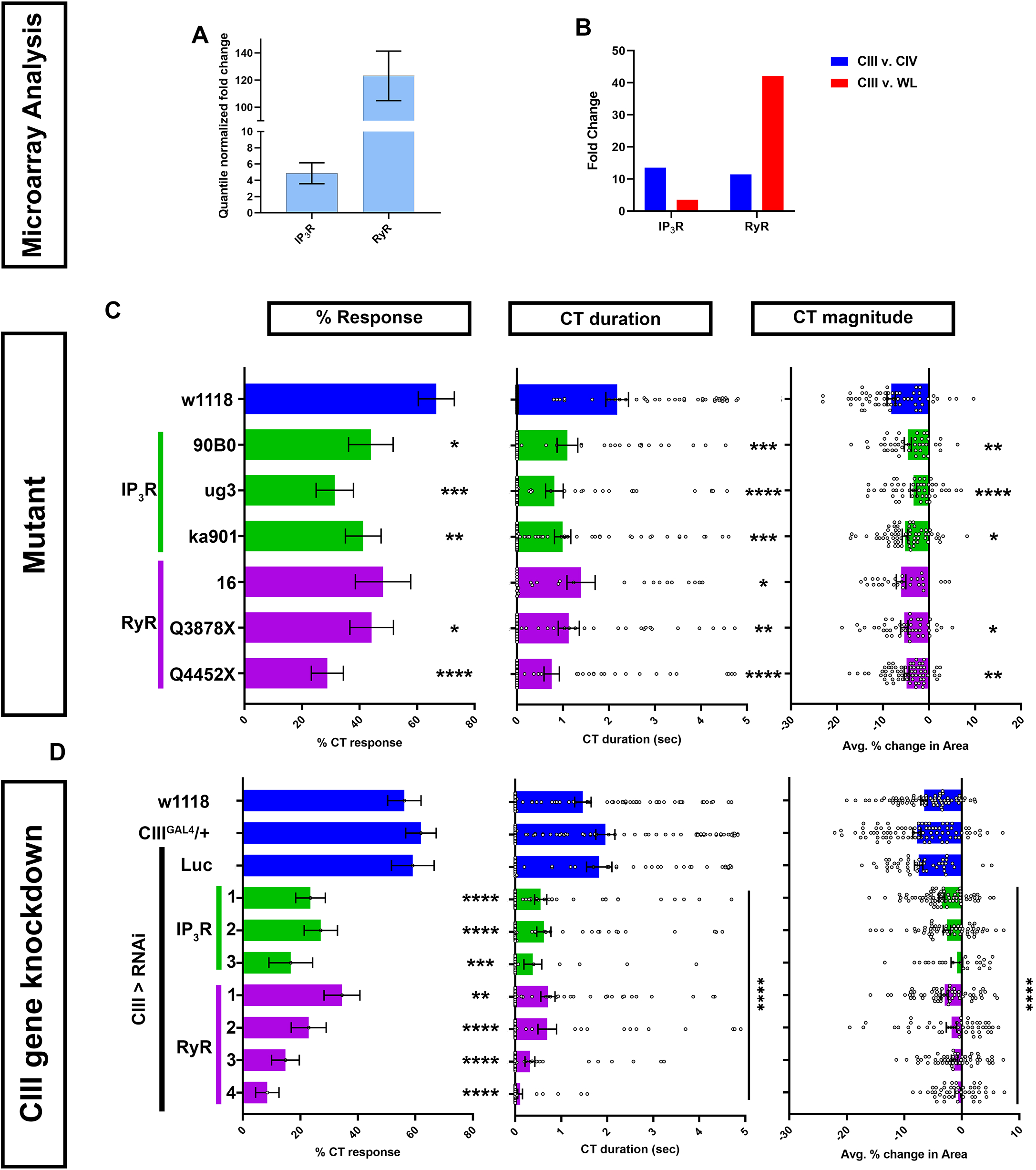
Ca^2+^ induced Ca^2+^ release genes IP_3_ and ryanodine receptors are required for cold-evoked behaviors. (**A-B**) Transcriptomic analysis using microarray datasets. (**A**) Quantile normalized expression of *RyR* and *IP*_*3*_*R* in CIII md neurons. (**B**) Comparative analysis of *IP*_*3*_*R* and *RyR* expression between CIII md neurons and CIV md neuron or whole larva (WL). (**C-D**) Cold plate analysis of RyR and IP_3_R. (**C**) *Drosophila* larvae heterozygous mutants for either *IP*_*3*_*R* (*ug3, ka901* and *90B0* alleles) or *RyR* (*16, Q3878X*, and *Y4452X* alleles). All statistical comparisons were made to *w1118*. N_Average_ = 49. We report %CT (left), CT duration in seconds (middle), CT magnitude (right). (**D**) *IP*_*3*_*R* or *RyR* knockdown specifically in CIII md neurons using *19-12*^*GAL4*^ driver. Control conditions include: *w1118, 19-12*^*GAL4*^*/+*, and *19-12>Luc*^*RNAi*^. We report %CT (left), CT duration in seconds (middle), CT magnitude (right). There is no statistical difference between the controls. Knockdown of *RyR* and *IP*_*3*_*R* was tested using multiple independent RNAi lines and comparisons made to *19-12*^*GAL4*^*/+*. N_Average_ = 55. Significant differences indicated via asterisks, where *p<0.05, **p<0.01, ***p<0.001, and ****p<0.0001.

Homozygous mutants for RyR or IP_3_R do not survive to the third instar larval developmental stage (Joshi et al., 2004) (data not shown). Therefore, we assessed cold-evoked behaviors of heterozygous mutants for IP_3_R including: IP_3_R^90B0^, a deficiency removing both IP_3_R and NMDAR genes, IP_3_R^ug3^, a missense mutant in the ligand binding domain that mutates polar serine to non-polar phenylalanine, and IP_3_R^ka901^, a mutation in the Ca^2+^ channel domain, where a conserved non-polar glycine is replaced with polar serine (Joshi et al., 2004; Venkatesh & Hasan, 1997). All three IP_3_R mutants had significant deficits in cold-evoked contraction behavior for %CT response, CT duration and magnitude compared to controls (**Figure 2C**). Three mutations for RyR were also tested for cold-evoked responses including RyR^16^, RyR^Q3878X^, and RyR^Y4452X^. RyR^16^ is a transposase mutant, where the translational start site and parts of the second exon are deleted. RyR^Q3878X^ is a nonsense mutant introducing a stop codon in the Ca^2+^ binding domain and RyR^Y4452X^ is another nonsense mutation, where the receptor is truncated at the transmembrane/pore domain (Gao et al., 2013). Heterozygous mutants for RyR^16^ had subtle, but statistically non-significant, reductions in cold-evoked %CT response and CT magnitude, however, CT duration was significantly reduced compared to control (**Figure 2C**). RyR^Q3878X^ and RyR^Y4452X^ heterozygous mutants both exhibited significant reductions in all three cold-evoked behavioral metrics (**Figure 2C**).

Mutant analyses revealed that IP_3_R and RyR whole animal heterozygotes showed significant reductions in cold-evoked CT behaviors. Next, we performed cell type specific gene knockdown for *RyR* and *IP*_*3*_*R* to test whether these genes functions in cold sensitive CIII md neurons. We tested multiple independent *UAS-RNAi* transgenes targeting both genes. RNAi-mediated gene knockdowns for *IP*_*3*_*R* and *RyR* led to severe reductions in cold-evoked CT behaviors, where %CT response, CT duration and magnitude were significantly lower than controls (**Figure 2D**). Impairments in cold-evoked CT behaviors for *IP*_*3*_*R* and *RyR* gene knockdowns were much stronger than mutants for the respective genes. Collectively, our genetic and behavioral studies implicate CICR molecular machinery including G_αq_, PLCβ, IP_3_R and RyR in cold nociceptive behavior.

Additionally, we sought to decipher whether genes required for proper cytosolic and ER Ca^2+^ homeostasis may play a role in *Drosophila* larval cold sensitive neurons for cold nociception. We assessed the roles of <u>s</u>tore <u>o</u>perated <u>C</u>a^2+^ <u>e</u>ntry (SOCE), and <u>S</u>arco/<u>e</u>ndoplasmic <u>r</u>eticulum <u>C</u>a^2+^ <u>A</u>TPase (SERCA), both of which are responsible for maintaining proper ER luminal Ca^2+^ levels, in cold-evoked behaviors. SOCE functions via multi-protein complex involving Stim, an ER luminal Ca^2+^ sensor, which serves as docking site between the plasma membrane (PM) and ER membrane, and Orai, a PM localized Ca^2+^ channel (Zheng et al., 2018). SERCA is a P-type Ca^2+^ transporter localized on the ER membrane that maintains proper ER luminal Ca^2+^ levels (Magyar et al., 1995). Firstly, transcriptomic analysis revealed that *SERCA* is enriched in CIII md neurons compared to CIV md neurons or whole larvae. *Stim* was highly enriched in CIII md neurons compared to CIV md neurons, however, only modestly enriched compared to whole larval transcriptome. *Orai* is only modestly enriched in CIII md neurons compared to CIV md neuron (**Supplementary Figure 3A**). CIII neuron specific knockdowns of *SERCA, Stim* or *Orai* lead to severe deficits in cold-evoked CT behaviors compared to controls (**Supplementary Figure 3B**). Collectively, our data reveal that GABA_B_ receptor, G protein signaling, CICR, and ER Ca^2+^ homeostasis are required in CIII md neurons for *Drosophila* larval cold nociceptive behavior.

### 3.3 GPCR and CICR signaling are not required for dendrite development or neural transmission

To test the possibilities that cold-evoked behavioral phenotypes of GPCR and CICR signaling impairments arise due to impaired neural development, neurotransmitter release, general excitability, and/or general stimulus processing, we performed morphometric as well as optogenetic analyses. We first assessed the effects of cell type specific gene knockdown in CIII md neurons on dendrite development. Morphologically CIII md neurons are characterized by having actin-rich dendritic filopodia emanating from major branches coupled to space-filling properties (Grueber et al., 2002). Cell-type specific knockdown of *GABA*_*B*_*-R2, IP*_*3*_*R* or *RyR* did not result in significant defects in CIII md neuron dendritic morphology compared to controls (**Figure 3A-C**). Sholl analysis, which is used for quantitatively assessing spatial branch distribution, did not reveal any significant changes from control (**Figure 3B**). We performed a principal components analysis on multiple morphological metrics (dendrite branch length, terminal branches, branch density, and sholl analysis) and we did not visualize any distinct clusters for gene knockdowns of *GABA*_*B*_*-R2, IP*_*3*_*R* or *RyR* compared to controls (**Figure 3C**).

**Figure 3:**
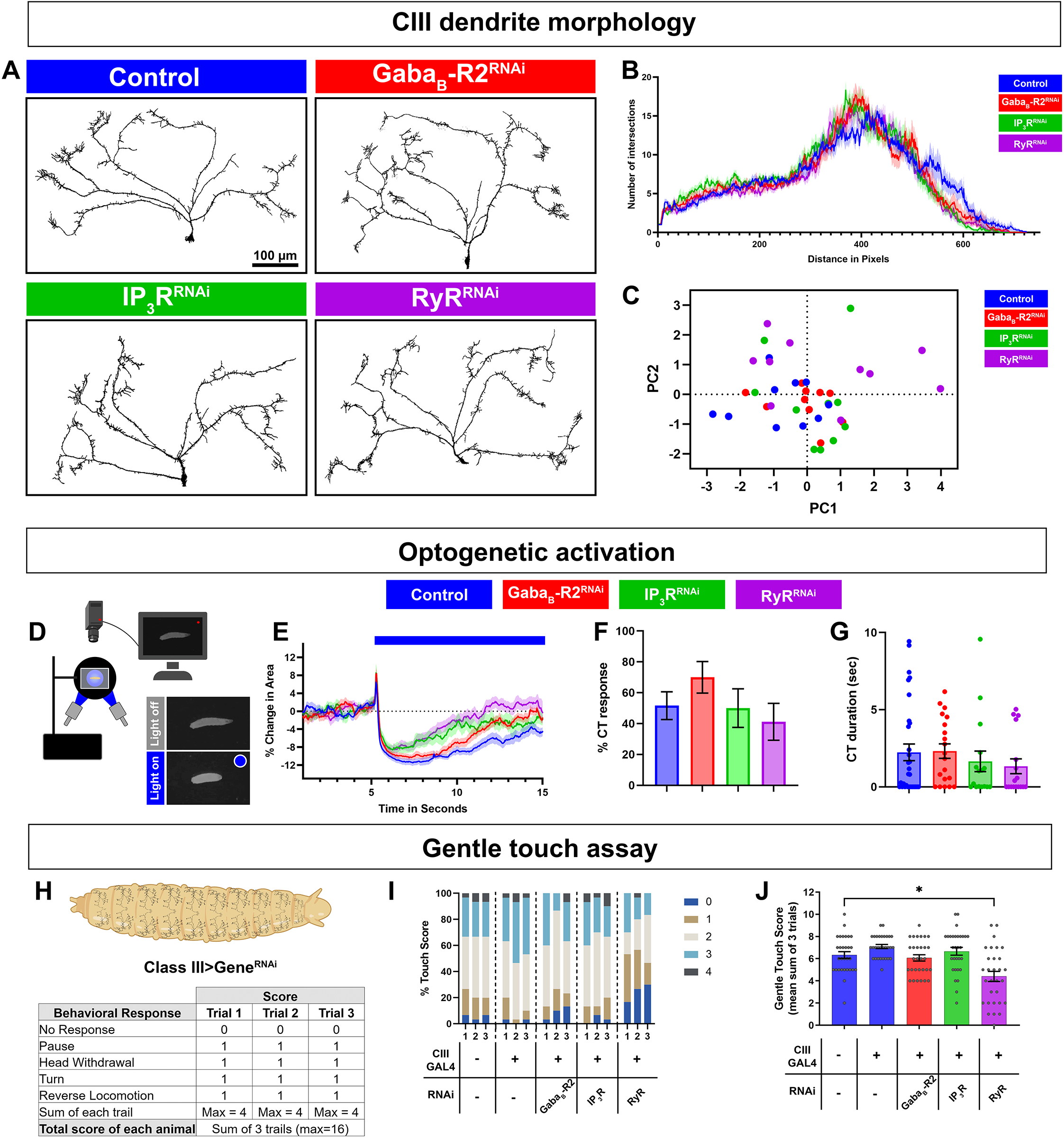
GPCR and CICR signaling are not required for CIII neuron dendrite development, general excitability, or neural transmission. (**A-C**) CIII md neuron dendrite morphometric analyses of cell type specific knockdown of *GABA*_*B*_-R2, *IP*_*3*_*R* or *RyR*. CIII md neurons were visualized using *nompC*^*GAL4*^*>GFP*. (**A**) Representative images of CIII md neuron (ddaF) dendrites with either *GABA*_*B*_-*R2, IP*_*3*_*R* or *RyR* knockdown compared to control (*Luc*^*RNAi*^). Scale bar = 100µm. (**B**) Sholl analysis of CIII md neuron dendrites, where numbers of intersections are plotted as a function of distance from the soma. (**C**) Principal components analysis of multiple morphometric parameters including total terminal branches, total dendritic length, branch density, max number of sholl intersections and distance at which the max intersections occur in Sholl analyses. N_Average_ = 10. (**D-G**) Neural activation of CIII md neurons using optogenetic actuator (ChETA) with cell type specific gene knockdown of *GABA*_*B*_*-R2, IP*_*3*_*R* or *RyR*. Genotype: *19-12>ChETA + gene-specific RNAi*. (**D**) Schematic of optogenetic stimulation apparatus. (**E**) Plot of change in area over time for *GABA*_*B*_*-R2, IP*_*3*_*R* or *RyR* knockdown compared to control (*Luc*^*RNAi*^). Blue bar represents 10 seconds of blue light exposure resulting in neural activation of CIII md neurons. (**F**) Optogenetic CIII md neuron activation evoked %CT response of *GABA*_*B*_*-R2, IP*_*3*_*R* or *RyR* knockdown compared to control. (**G**) CT duration in seconds for *GABA*_*B*_*-R2, IP*_*3*_*R* or *RyR* knockdown compared to control. (**E-G**) N_Average_ = 20. Statistics: %CT response: Fisher’s exact with Benjamini-Hochberg for multiple comparison. CT duration: One-way ANOVA with Holm-Šídák’s for multiple comparisons. (**H-J**) *Drosophila* larval responses to innocuous mechanical stimulation. (**H**) Gentle touch assay scoring schematic. Each behavioral response receives 1 point and no points, if there was no response, for a maximum of 4 points per trial. (**I**) Proportion of each score per trial for each genotype. (**J**) Average total touch scores for each larva per genotype. N_Average_ = 30. Statistics: Kruskal-Wallis with Benjamini, Krieger and Yekutieli for multiple comparisons. Significant differences indicated via asterisks, where *p<0.05, **p<0.01, ***p<0.001, and ****p<0.0001.

Deficits in cold-evoked behavioral responses could be a byproduct of the inability of CIII md neurons to defects in general excitability and/or defects in properly communicating with downstream neurons due to lack of neurotransmission or action potential propagation. To assess neural excitability and neurotransmitter release, we co-expressed ChETA, an excitatory optogenetic channel, with gene knockdown for *GABA*_*B*_*-R2, IP*_*3*_*R* or *RyR* in CIII md neurons (**Figure 3D**).

*Drosophila* larvae knocked down for *GABA*_*B*_*-R2, IP*_*3*_*R* or *RyR* in CIII md neurons were still able to perform CT behavior upon optogenetic activation suggesting that CIII md neurons are still able to generate action potentials and release neurotransmitter to downstream neurons (**Figure 3E**). Initial CT magnitude and %CT response between CIII md neuron specific gene knockdown and control were not significantly different (**Figure 3F-G**). These data indicate that general CIII md neuron function is not dependent on expression of GPCR and CICR genes.

### 3.4 GABAB-R2 and IP3R function in stimulus discrimination but RyR is required for multimodal processing

CIII md neurons are multimodal sensor and respond to both noxious cold and innocuous mechanical stimuli. Thus far our data indicate that GPCR and CICR genes are required for cold nociception but are not responsible for dendrite development or general excitability. We hypothesize that the ability of CIII md neurons to detect and respond to innocuous mechanical stimuli is not dependent on GPCR and CICR signaling. To assess innocuous mechanical sensitivity of *Drosophila* larvae, we performed gentle touch assays using a fine brush bristle on animals with CIII md neuron specific gene knockdowns for *GABA*_*B*_*-R2, IP*_*3*_*R* or *RyR* (Kernan et al., 1994). Responses of *Drosophila* larvae to innocuous mechanical stimulation include pause, head withdrawal, turn and/or reverse locomotion, where an animal receives a score for performing each behavior, max score of 4 for each trail (**Figure 3H**) (Himmel, Sakurai, et al., 2021; Kernan et al., 1994; Turner et al., 2016). Gentle touch-evoked responses of *GABA*_*B*_*-R2* and *IP*_*3*_*R* knockdown in CIII md neurons were not significantly different from controls (**Figure 3I-J**). In contrast, *RyR* knockdown led to significant decreases in total gentle touch score compared to controls. Assessing individual trial data for proportions of gentle touch score revealed a greater portion of *RyR* knockdown larvae having no response (NR) and reduced proportion of animals performing a greater number of behaviors (Touch score 2 or higher) (**Figure 3I-J**). Our data reveal that *GABA*_*B*_*-R2* and *IP*_*3*_*R* are not required for innocuous mechanical sensation, whereas *RyR* knockdown leads to deficits in innocuous mechanosensation. Collectively, these findings indicate that *GABA*_*B*_*-R2* and *IP*_*3*_*R* function in noxious cold stimulus discrimination, whereas *RyR* has a multimodal requirement for cold and innocuous touch sensation.

### 3.5 GPCR and CICR signaling are required for cold-evoked Ca2+ dynamics

CIII md neuron multimodality arises at least in part due to differential Ca^2+^ responses to stimuli, where high intracellular Ca^2+^ is observed in response to noxious cold and relatively low intracellular Ca^2+^ levels in response to innocuous touch. We hypothesize that GABA_B_ and CICR signaling mechanisms are required for cold-evoked high intracellular Ca^2+^ levels in CIII neurons. To evaluate Ca^2+^ responses, we utilized a transient Ca^2+^ sensor GCaMP6. Using a cell type specific *GAL4* driver, we expressed GCaMP6 and RNAi for either *GABA*_*B*_*-R2, IP*_*3*_*R* or *RyR*, in CIII neurons and imaged *in vivo* Ca^2+^ responses in intact *Drosophila* larvae (**Figure 4A**). In controls, there is a robust increase in GCaMP6 fluorescence in CIII md neuron cell bodies in response to noxious cold. Upon *GABA*_*B*_*-R2* knockdown, there is a nearly 35% reduction in cold-evoked Ca^2+^ transient compared to controls. *IP*_*3*_*R* or *RyR* knockdown in CIII md neurons leads to nearly 50% reduction Ca^2+^ response compared to controls (**Figure 4B-C**). Therefore, cold-evoked intracellular increases Ca^2+^ are dependent on GPCR and CICR signaling in CIII md neurons.

**Figure 4:**
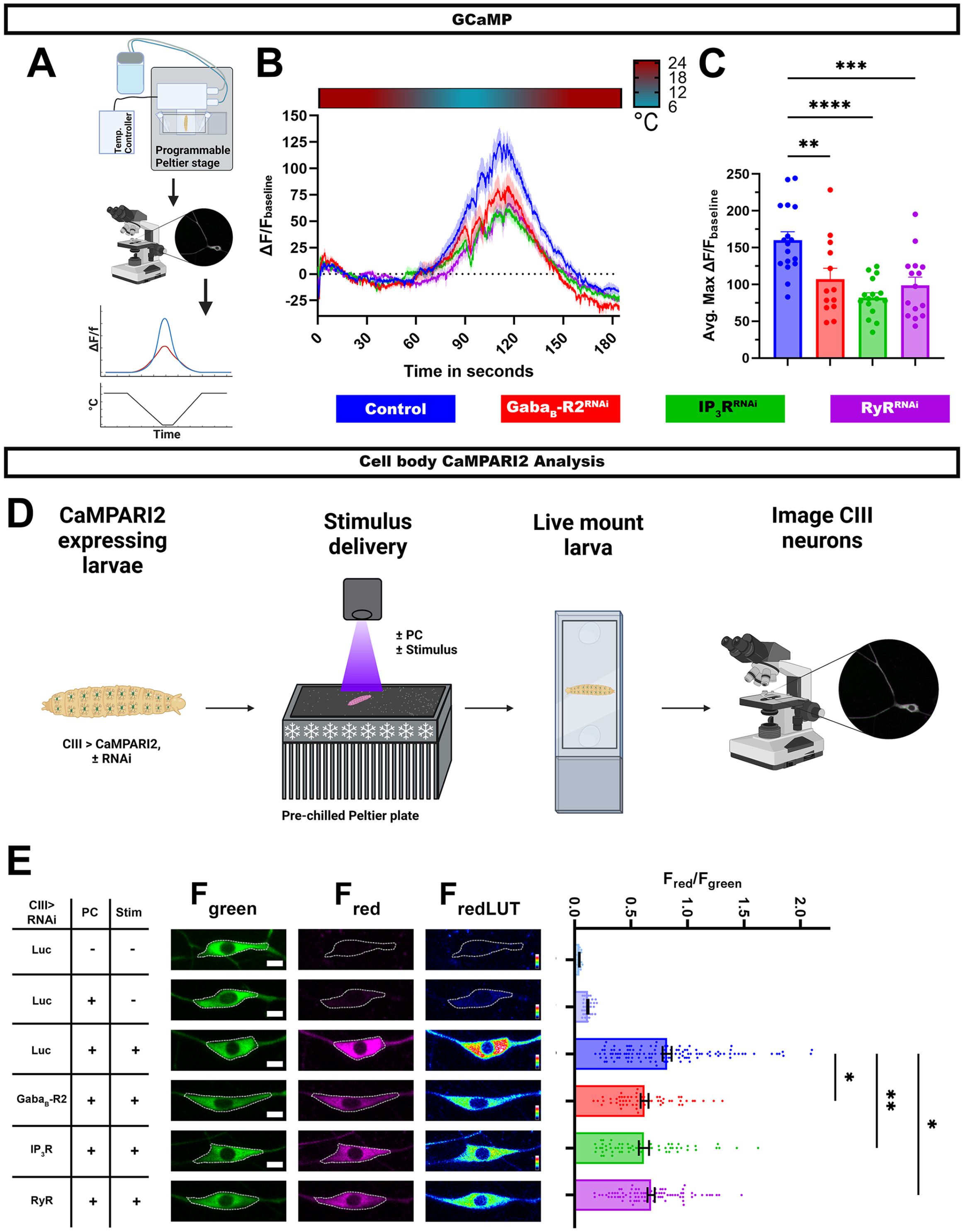
GPCR and CICR signaling are required for cold-evoked Ca^2+^ dynamics. (**A-C**) Cold-evoked *in vivo* GCaMP6 imaging of CIII md neuron cell bodies. (**A**) Schematic of cold stimulus delivery and imaging system. (**B**) Plot of ΔF/F_baseline_ over time of CIII md neuron cell body (ddaF) expressing GCaMP6m and gene knockdown for *GABA*_*B*_*-R2, IP*_*3*_*R*, or *RyR* compared to control (*Luc*^*RNAi*^). (**C**) Max ΔF/F_baseline_ for cold-evoked GCaMP response in CIII md neuron cell body. N_Average_ = 15. Statistics: One-way ANOVA with Holm-Šídák’s for multiple comparisons. (**D-E**) *in vivo* analysis of CaMPARI2 responses to noxious cold in CIII md neurons expressing gene knockdowns for *GABA*_*B*_*-R2, IP*_*3*_*R*, or *RyR* compared to control (*Luc*^*RNAi*^). (**D**) Graphical schematic of cold plate assay combined with photoconverting (PC) light and post hoc imaging. (**E**) Left: Representative images of CIII md neurons expressing CaMPARI2 and gene specific *RNAi*. Cell bodies are outline in dashed white line. Scale bar: 5µM. Right: CaMPARI2 responses of CIII md neurons (ddaA and ddaF) plotted as fluorescence ratios of F_red_/F_green_. N_Average_ = 62. Statistics: Kruskal-Wallis with Benjamini, Krieger and Yekutieli for multiple comparisons. Significant differences indicated via asterisks, where *p<0.05, **p<0.01, ***p<0.001, and ****p<0.0001.

Assessing Ca^2+^ transient using GCaMP6 allows for time resolved Ca^2+^ responses to stimulus, however, this experimental design has restrictions on spatial resolution and depend on restraining the animal during stimulus delivery. To overcome these experimental design limitations, we visualized Ca^2+^ responses via a Ca^2+^ integrator, CaMPARI2, which allows for greater spatial resolution and larvae can be unrestrained and freely moving during the stimulus delivery (Moeyaert et al., 2018).

We co-expressed *CaMPARI2* and *GABA*_*B*_*-R2, IP*_*3*_*R* or *RyR* RNAi in CIII md neurons and assessed cold-evoked CaMPARI2 responses along the body wall (**Figure 4D**). CaMPARI2’s fluorescence changes from green to red in presence of high intracellular Ca^2+^ and photoconverting (PC) light, where neuronal responses to stimuli are reported as F_red_/F_green_ ratios (Fosque et al., 2015). We controlled for exposure to PC light, stimulus, and expression of the RNAi construct. When analysing neuronal cell bodies, control CIII md neurons have significantly higher F_red_/F_green_ ratios compared to two no stimulus controls, one with or without PC light (**Figure 4E**). Knockdowns of *GABA*_*B*_*-R2, IP*_*3*_*R* or *RyR* in CIII md neurons results in significantly lower cold-evoked CaMPARI2 responses compared to controls (**Figure 4E**).

In addition to analyzing CaMPARI2 responses at the cell body, we took advantage of greater spatial resolution of CaMPARI2 to analyze F_red_/F_green_ ratios along the dendritic arbor. We implemented general principles of Sholl analysis to analyze neuronal CaMPARI2 red and green fluorescence intensities along the dendrite (**Supplementary Figure 4A**). A similar approach has been utilized to analyze the neuronal cytoskeleton (Nanda et al., 2021), however, we created new higher throughput custom FIJI scripts to analyze CaMPARI2 in somatosensory neurons. As with the cell body analysis, cold-evoked CaMPARI2 responses along the dendrites were also reduced for *GABA*_*B*_*-R2, IP*_*3*_*R* or *RyR* knockdowns (**Supplementary Figure 4C**). The observed reductions in F_red_/F_green_ ratios are not due to differences in dendritic area of CIII md neurons across genotypes (**Supplementary Figure 4B**). Collectively, these results demonstrate that GABA_B_ and CICR genes are required for cold-evoked Ca^2+^ responses of CIII md neurons.

### 3.6 GABA_B_-R2 and ryanodine receptors are required for bursting in cold sensing neurons

Here, we have demonstrated that GABA_B_-R2 and ryanodine receptors are required for cold-evoked behaviors, where they mediate increases in cytosolic Ca^2+^ in CIII md neurons in response to noxious cold temperatures. We hypothesized that GABA_B_-R2 and RyR are required for proper cold-evoked CIII firing patterns. We performed extra-cellular recordings of CIII neurons in third instar larval filets. We exposed *Drosophila* larvae to three different stimuli (20°C, 15°C and 10°C), where there is a rapid ramp down to the target temperature and followed with continuous exposure to the stimulus temperature, for a total of 60 seconds (**Figure 5A**). In controls, a rapid decrease in temperature causes increased firing composed mainly of bursts, defined as three or more consecutive spikes with an inter-spike interval of less than 0.15 seconds (Himmel, Letcher, et al., 2021; Himmel, Sakurai, et al., 2021) (**Figure 5A, C**). During steady state cold exposure, CIII md neurons generally show tonic spiking (Himmel, Letcher, et al., 2021) (**Figure 5A, D**). First, we assessed how loss of GABA_B_-R2 receptor affected cold-evoked firing pattern of CIII md neurons (**Figure 5A-B**). *GABA*_*B*_*-R2* knockdown led to significant reductions in average spike rate at room temperature (pre-stimulus baseline) and at 20°C or 15°C cold exposure compared to controls (**Figure 5A-B**). However, we did not observe any change in cold-evoked firing upon 10°C exposure (**Figure 5A-B**). To further characterize how *GABA*_*B*_*-R2* knockdown effects cold-evoked firing, we examined the composition of bursting and tonic activity in 10 second bins for the 10°C stimulus. *GABA*_*B*_*-R2* knockdown led to a disorganized bursting pattern in cold exposed CIII md neuron, where on average there was a 28% increase in burst firing rate compared to controls (**Figure 5C, E**). Meanwhile, there were significant changes in tonic activity with an average of 28% reduction in tonic firing rate in *GABA*_*B*_*-R2* depleted neurons compared to controls (**Figure 5D, E**). GABA_B_-R2 signaling is required for proper cold-evoked electrical activity in CIII md neurons.

**Figure 5:**
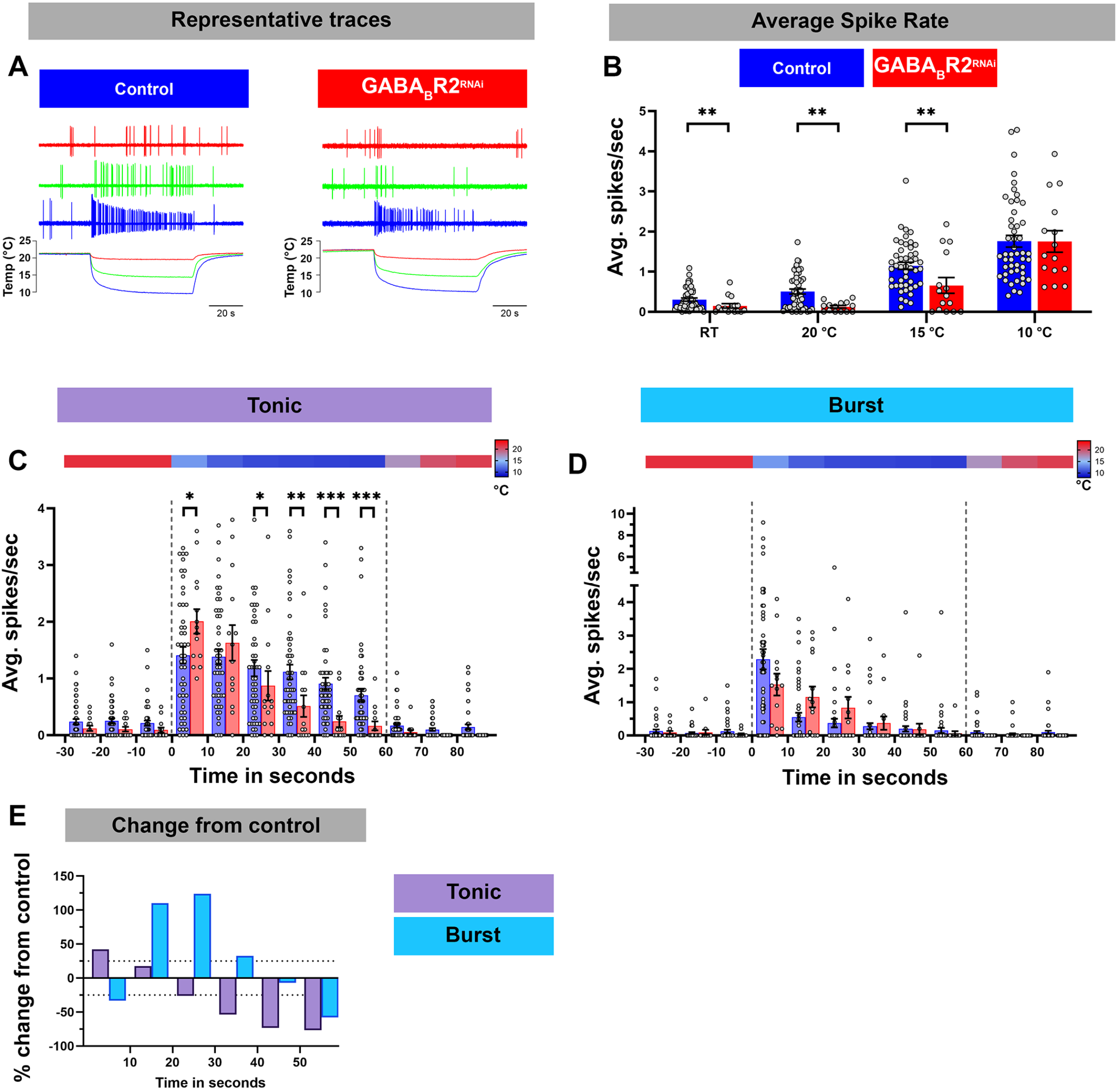
GABA_B_-R2 receptor is required for proper composition of cold-evoked tonic and bursting pattern in CIII md neurons. (**A**) Representative physiological firing traces of CIII md neurons with *GABA*_*B*_*-R2* knockdown compared to controls during each stimulus temperature (20°C, 15°C, 10°C) and below shows real time temperature exposure regimes of *Drosophila* larval filet. (**B**) Average spikes/seconds of CIII md neurons at room temperature (RT) or cold stimulations (20°C, 15°C, 10°C). N_Average_ = 32. Statistics: Multiple Mann-Whitney test with Benjamini, Krieger and Yekutieli for multiple comparisons. (**C-D**) 10°C stimulus evoked tonic spike rate (**C**) and bursting spike rate (**D**). Individual spikes are considered to bursts, if inter spike interval is less than 0.15 seconds for at least 3 spikes and rest of the spikes counted as tonic spikes. Heatmap above each graph represents stimulus temperature. Spike rate is calculated in 10 second bins. Statistics: Multiple Mann-Whitney tests with Benjamini, Krieger and Yekutieli for multiple comparisons. (**E**) Percent change from control for bursting or tonic firing in GABA_B_-R2 knockdown. Significant differences indicated via asterisks, where *p<0.05, **p<0.01, and ***p<0.001.

Next, we assessed how loss of RyR signaling affected the cold-evoked firing pattern of CIII md neurons (**Figure 6A-C, G**). *RyR* knockdown led to a significant decrease in average spike rate at room temperature (pre-stimulus baseline) and at 20°C or 15°C cold exposure compared to their respective controls (**Figure 6A, Supplementary Figure 5A**). Similar to our findings with GABA_B_-R2, the average spike rate was not significantly different in *RyR* depleted neurons relative to controls in response to a 10°C stimulus (**Figure 6A**). To further characterize the effects on *RyR* depletion, we parsed out bursting and tonic spikes. *RyR* knockdown led to significant reductions, an average of 55% decrease, in cold-evoked bursts over the stimulus window (**Figure 6B, G**). Interestingly, *RyR* knockdown increased the tonic firing rate resulting in an average 53% percent increase in tonic firing rate within the first 30 seconds of stimulus compared to controls (**Figure 6C, G**). During last 30 seconds of 10°C stimulus, there was an average 27% increase in tonic firing rate compared to controls (**Figure 6C, G**). Together, our physiological analyses of CIII neuron specific *RyR* knockdown revealed that RyR functions to promote bursting and restrict tonic firing in CIII md neurons.

**Figure 6:**
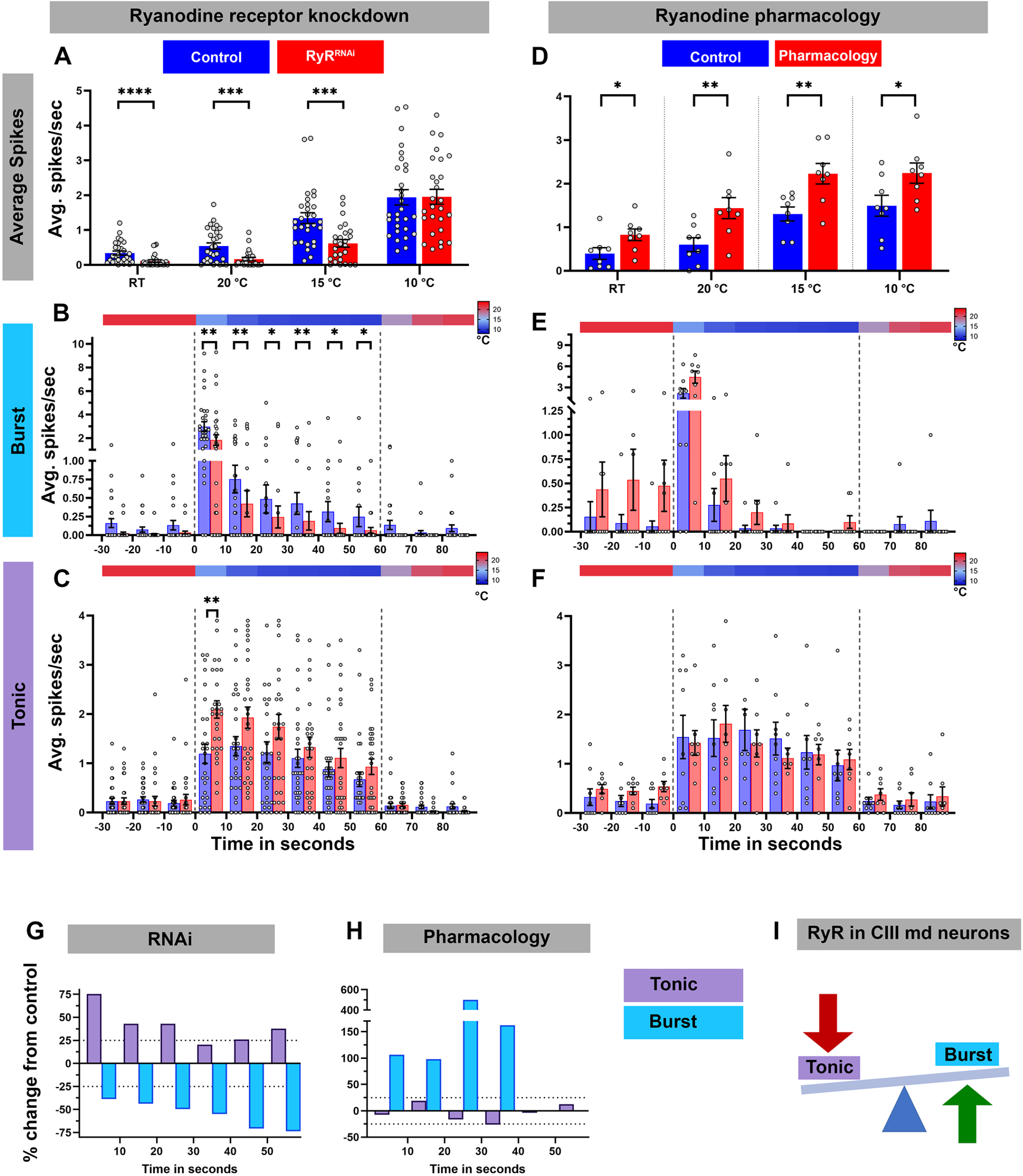
Ryanodine receptor mediates cold-evoked bursting in CIII md neurons. (**A-C**) Average spike rate of GFP tagged CIII md neurons expressing *RyR*^*RNAi*^ compared to control. (**A**) Average spikes/seconds of CIII md neurons at room temperature (RT) or cold stimulations (20°C, 15°C, 10°C). N_Average_ = 29. Statistics: Multiple Mann-Whitney tests with Benjamini, Krieger and Yekutieli for multiple comparisons. (**B-C**) 10°C stimulus evoked bursting spike rate (**B**) and tonic spike rate (**C**). Individual spikes are considered to bursts, if inter spike interval is less than 0.15 seconds for at least 3 spikes and rest of the spikes counted as tonic spikes. Heatmap above each graph represents stimulus temperature. Spike rate is calculated in 10 second bins. Statistics: Multiple Mann-Whitney test with Benjamini, Krieger and Yekutieli for multiple comparisons. (**D-F**) Average spike rate of GFP tagged CIII md neurons in response to cold stimulus under ryanodine (10^−6^ M) pharmacology application. (**D**) Average spikes/seconds of CIII md neurons at room temperature (RT) or cold stimulations (20°C, 15°C, 10°C). N_Average_ = 8. Statistics: Multiple Wilcoxon tests with Benjamini, Krieger and Yekutieli for multiple comparisons. (**E-F**) 10°C stimulus evoked bursting spike rate (**E**) and tonic spike rate (**F**). (**G-H**) Percent change from control for bursting or tonic firing in RyR knockdown (**G**) or ryanodine application (**H**) at 10°C. (**I**) CICR mechanism through RyR functions in generating cold-evoked burst firing pattern in CIII md neuron. Significant differences indicated via asterisks, where *p<0.05, **p<0.01, ***p<0.001, and ****p<0.0001.

To independently investigate the relative effects of *RyR* depletion, we performed pharmacological experiments involving bath application of ryanodine (10^−6^ M) to larval filet preparations to activate RyRs, and assessed cold-evoked firing patterns of CIII md neurons (**Figure 6D-F, H, Supplementary Figure 5B**). In contrast to our observations with *RyR* knockdown **(Figure 6A)**, ryanodine application led to significant increases in average firing rate at room temperature, as well as during 20°C, 15°C or 10°C stimulus (**Figure 6D, Supplementary Figure 5B**). We next explored the role of ryanodine application on the types of firing patterns in CIII md neurons and did not find statistically significant differences, however, percent change from control analyses revealed an increase in bursting upon ryanodine application. In bath-applied ryanodine conditions, CIII md neurons exposed to 10°C exhibited an average 144% increase in bursting activity compared to controls (**Figure 6E, H)**. Overall, ryanodine application did not affect the tonic firing rate (**Figure 6F, H**). These data support a role for RyR in promoting cold-evoked bursting activity in CIII md neuron (**Figure 6I**).

### 3.7 TRP channel Pkd2 and IP3R genetically interact in cold nociception

*Drosophila* larval cold-evoked behaviors are dependent on TRP channels, specifically Pkd2, and cold-evoked increases in CIII md neuron cytosolic Ca^2+^ levels require Pkd2 (Turner et al., 2016). CICR mechanisms function in amplifying second messenger Ca^2+^ signaling, which relies upon Ca^2+^ influx through pathways including TRP channel mediated increases in cytosolic Ca^2+^. We hypothesize that Pkd2 and IP_3_R function synergistically in sensing noxious cold stimuli. To assess whether, *Pkd2* and *IP*_*3*_*R* genetically interact, we utilized a transheterozygote mutant approach where there is a single mutant copy for each gene. Single allele mutants for either *Pkd2*^*1*^ or *IP*_*3*_*R*^*ka901*^ led to significant reductions in cold-evoked %CT response, CT duration and CT magnitude compared to controls (**Figure 7**). *Drosophila* larvae transheterozygous mutant for *Pkd2*^*1*^ and *IP*_*3*_*R*^*ka901*^ had more severe reductions in cold-evoked CT behavior compared to either single allele mutants or controls (**Figure 7**). These analyses demonstrate a synergistic genetic interaction between *IP*_*3*_*R* and *Pkd2* suggesting TRP channel and CICR signaling may operate together to mediate cold-evoked nociceptive behavior.

**Figure 7:**
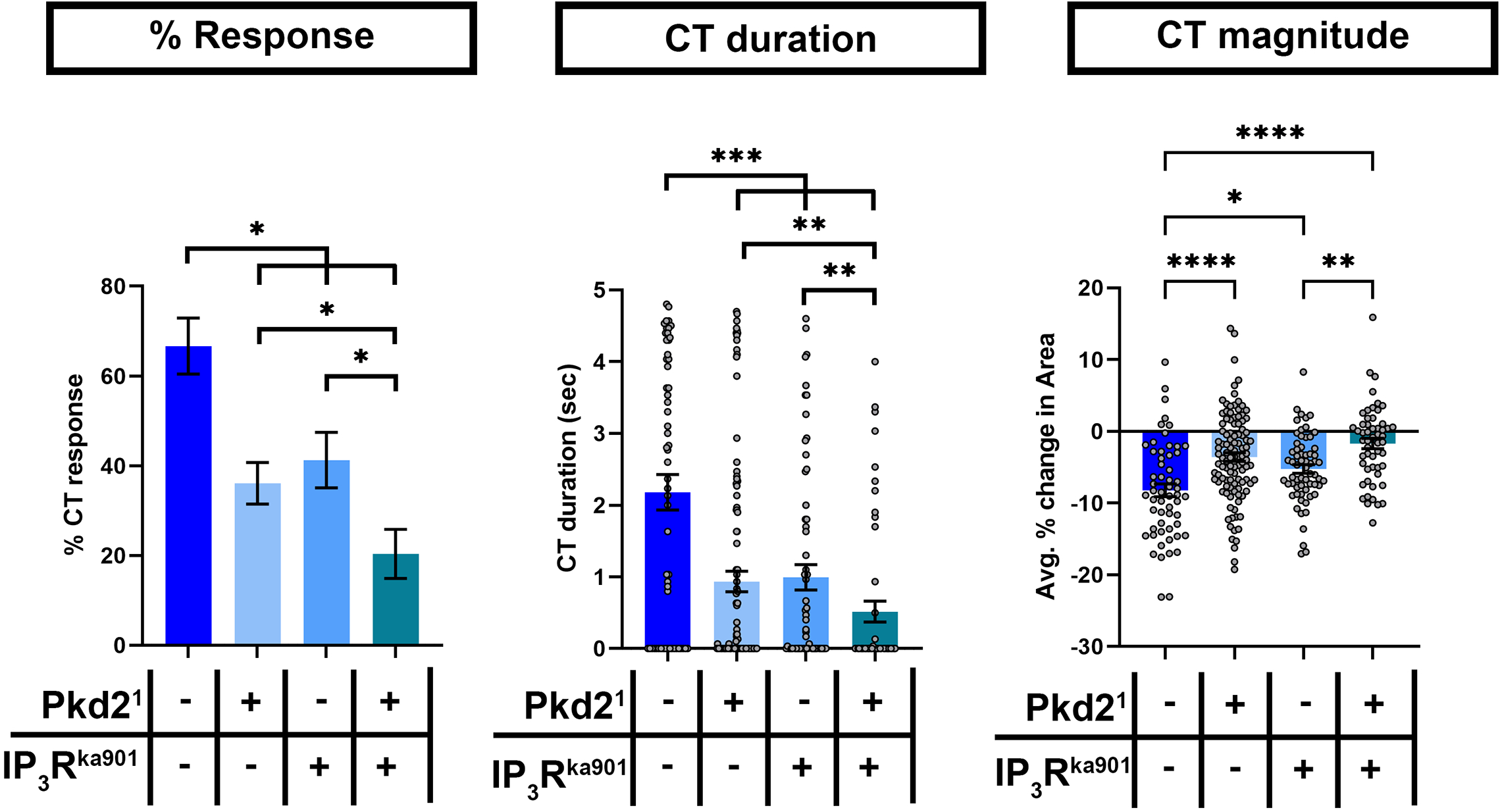
Pkd2 and IP_3_R genetically interact in cold nociception. Cold plate analysis of *Drosophila* larvae mutant for either/both *Pkd2*^*1*^ and *IP*_*3*_*R*^*ka901*^. We report %CT (left), CT duration in seconds (middle), CT magnitude (right). Genotypes include: *w1118, Pkd2*^*1*^*/+*, and *IP*_*3*_*R*^*ka901*^*/+, Pkd2*^*1*^*/IP*_*3*_*R*^*ka901*^. N_Average_ = 70. Significant differences indicated via asterisks, where *p<0.05, **p<0.01, ***p<0.001, and ****p<0.0001.

## 4 Discussion

We have demonstrated that both metabotropic GABA_B_ receptor and CICR signaling pathways are required in *Drosophila melanogaster* larva for cold nociceptive behaviors (**Figure 8**). First, we have shown that *Drosophila* larvae mutant for *GABA*_*B*_, *IP*_*3*_*R*, or *RyR* have impaired cold-evoked CT responses. GABA_B_, IP_3_R and RyR function is specifically required in CIII md neurons for cold nociception, where gene knockdown results in reductions of cold-evoked Ca^2+^ responses. Additionally, our extracellular recordings show that cold-evoked CIII md neuronal bursting response is mediated by RyR presumptively related to Ca^2+^ release from ER stores. Behavioral and functional impairments in cold nociception are not due to structural or developmental defects of CIII md - between *GABA*_*B*_*-R2, IP*_*3*_*R* or *RyR* knockdown and controls. Furthermore, loss of *GABA*_*B*_*-R2, IP*_*3*_*R* or *RyR* did not lead to significant reductions in CT responses when CIII md neurons were optogenetically stimulated indicating that CIII-mediated neurotransmission is functional.

**Figure 8:**
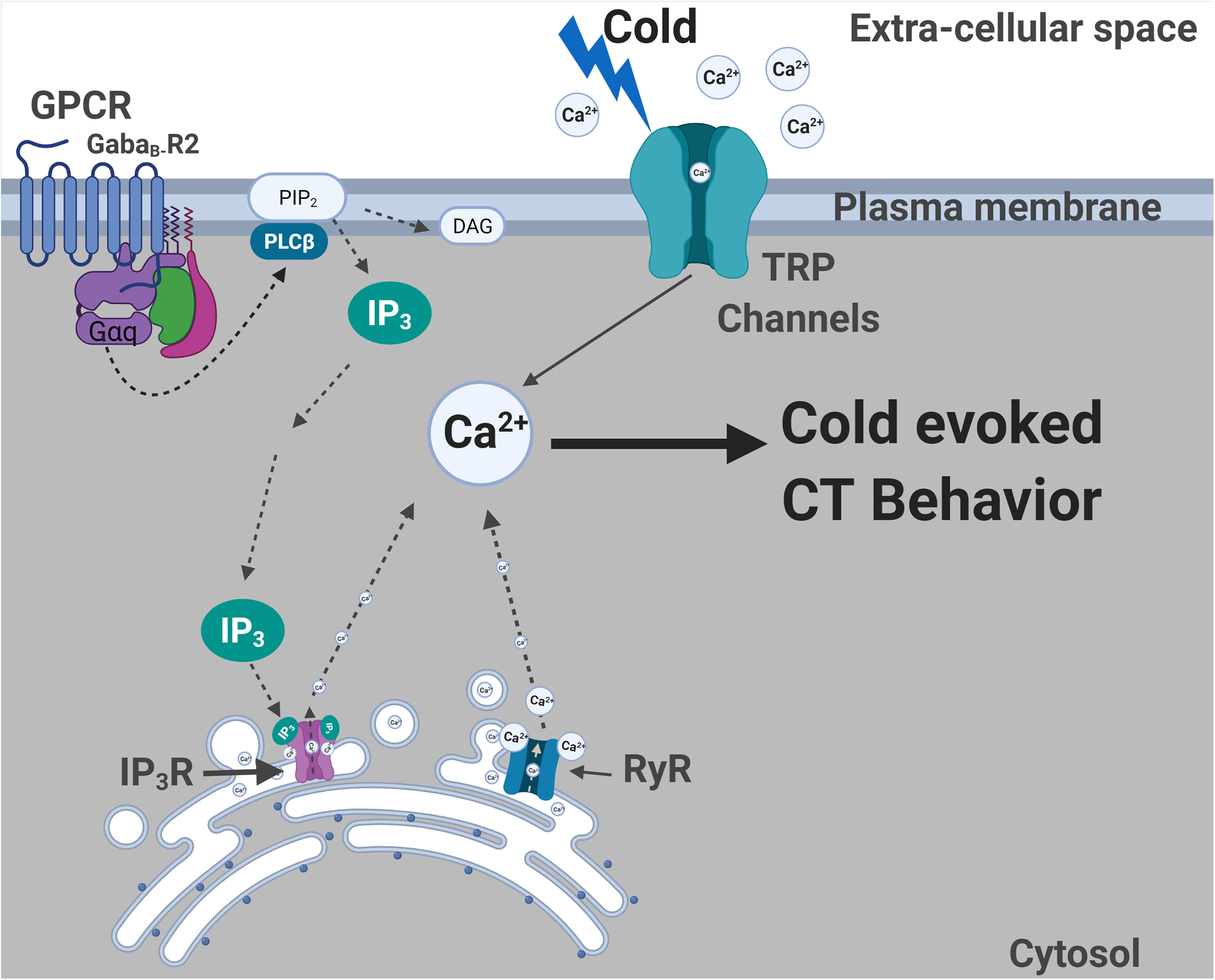
Summary of molecules required for cold nociceptive behaviors. Plasma membrane localized TRP channels (Turner et al., 2016) and GPCR (GABA_B_-R2) are required for cold-evoked behaviors. G_αq_ and Plc signalling is also required for cold nociception. ER localized IP_3_R and RyR channels are required for cold-evoked CT response. GABA_B_-R2, IP_3_R and RyR are all required for cold-evoked Ca^2+^increases.

We show that GABA_B_-R2 in conjunction with G_αq_/PLC mediated GPCR signaling is required for cold-evoked CT responses and evoked increases in cytosolic Ca^2+^. *GABA*_*B*_*-R2/IP*_*3*_*R* signaling are required for cold-evoked neural responses in CIII md neurons, when assessed by GCaMP or CaMPARI. *In vivo* GCaMP analyses indicated that cold-evoked Ca^2+^ increases at the cell body are reduced during temperature ramp down and return to baseline in *GABA*_*B*_*-R2/IP*_*3*_*R* knockdowns compared to control. Additionally, CaMPARI analyses reveal that these reductions in cold-evoked Ca^2+^ are not restricted to the cell body but also seen on dendrites. Our results indicate that one GPCR downstream signaling mechanism via G_αq_, PLCs and IP_3_R is required for cold-evoked behaviors. However, GPCR activation of G_αq_ and PLCs leading to local depletion of PIP_2_ generating IP_3_ and DAG may have other possible impacts on plasma membrane localized ion channels. PIP_2_ positively enhances conductance of voltage-gated Ca^2+^ channels (VGCCs), where PIP_2_ depletion through activation of either voltage sensitive lipid phosphatases, PLCs and/or G_q_ signaling pathways leads to inhibition of select VGCCs (Heneghan et al., 2009; Kammermeier et al., 2000; Liu et al., 2006; Suh et al., 2012). In rodents, dorsal root ganglion (DRG) neuronal pre-synaptic GABA_B_ receptor function regulates N-type Ca^2+^ currents in a voltage dependent and independent manner (Callaghan et al., 2008; Li et al., 2002; Raingo et al., 2007; Richman & Diverse-Pierluissi, 2004). Metabotropic GABA receptors function in neuroprotection, where research in cerebellar granule cell cultures has shown that GABA_B_ mediated activation of RhoA and Rac functions through G_13_ to promote synaptic activity and reduce apoptotic cascades (Wang et al., 2021). In neonatal hippocampal cultures, GABA_B_ receptors have been shown to colocalize with G_αq_ and function through IP_3_R and PKCα enhancing L-type Ca^2+^ current (Karls & Mynlieff, 2015). Additionally, GABA_B_ receptors are required in olfactory sensory neuron for *Drosophila* larval feeding responses (Slankster et al., 2020). These recent findings, including our reported findings, support new emerging roles for GABA_B_ receptors in signaling pathways promoting stimulus evoked behaviors, where GABA_B_ receptors are required for neural activity.

It is as yet unclear how metabotropic GABA receptors are activated upon cold exposure. Potentially, GABA, the endogenous ligand, might be functioning at axon terminals of cold sensitive somatosensory neurons. Alternatively, *Drosophila* larval somatosensory neuronal cell bodies are wrapped in two types of glial cells (perineurial and sub-perineurial) (Yadav et al., 2019). Glial-to-neuron communications in CIII md neurons may lead to cold-evoked activation of GABA_B_ receptors. In another scenario, the activation of GABA_B_ receptors may function via an as yet unidentified ligand, which may be released upon cold exposure locally near somatosensory neurons. The TrpA1 channel is sensitive to both noxious heat and chemical stimuli (Arenas et al., 2017). A potential ligand for TrpA1 activation was identified as reactive oxygen species (ROS) and both *D. melanogaster* and *S. mediterranea* species exhibited stimulus evoked increases in ROS leading to TrpA1 activation (Arenas et al., 2017). Identifying potential physiological ligands that may be generated upon cold exposure may provide insights into requirements of new signaling mechanisms in nociception and are fertile ground for future investigations.

ER Ca^2+^ homeostasis in CIII md neurons is necessary for nociceptive behavioral responses. Our results indicate that ER Ca^2+^ release (IP_3_R & RyR) and reuptake channels (SERCA, STIM, & ORAI) are required in CIII md neurons for cold-evoked CT responses. Expectedly, CIII neuron specific impairments in ER Ca^2+^ release channels led to significant reductions in cold-evoked increases in cytosolic Ca^2+^ which may ultimately disrupt electrical activation patterns and/or neurotransmission. Ryanodine receptor’s primary ligand is Ca^2+^ and the likely source of Ca^2+^ is through cold sensitive Ca^2+^ permeant channels (Woll & Van Petegem, 2022). However, IP_3_R mediated Ca^2+^ release requires either IP_3_ production through G-protein signaling or cytosolic Ca^2+^ binding to the receptor (Woll & Van Petegem, 2022). IP_3_R and RyR have a biphasic sensitivity to Ca^2+^ binding and precise cytosolic Ca^2+^ required for channel activation/inactivation are dependent on specific cell types and receptor isoforms (Bezprozvanny et al., 1991; Laver, 2018). Interestingly, GABA_B_-R2 and IP_3_R, which are known to function in the same molecular pathway in the neonatal hippocampus (Karls & Mynlieff, 2015), are required for cold nociception and dispensable for innocuous mechanosensation, however, RyR is required for innocuous mechanosensation and noxious cold detection (Karls & Mynlieff, 2015). This suggests that Ca^2+^ sensitivity curves for both RyR and IP_3_R may be functionally different in CIII md neurons to allow for gating innocuous/noxious responses.

In response to noxious cold, CIII md neurons exhibit significant increases in cytosolic Ca^2+^ (Turner et al., 2016), coupled with temperature-dependent increases in electrical activity (Himmel, Letcher et al., 2021) where the CIII electrical responses to cold stimulus are linearly encoded as stimulus intensity increases. There are also linear increases in cold-evoked CT behavioral responses to increases in cold stimulus or optogenetic mediated activation of CIII md neurons (Turner et al., 2016). *GABA*_*B*_*-R2* or *RyR* knockdown resulted in reduced cold sensitivity, where in the innocuous cool range *GABA*_*B*_*-R2 or RyR* depleted neurons require a greater magnitude cold stimulus to achieve similar spike rates relative to controls (*i*.*e. GABA*_*B*_*-R2 or RyR* knockdown at 15°C vs. Control at 20°C have spike rates that show no statistical difference; **Figure 5B, 6A**). However, in the noxious range (≤10°C), we did not observe significant differences in spike rates for *GABA*_*B*_*-R2 or RyR* depleted neurons relative to controls suggesting that *GABA*_*B*_*-R2 or* RyR have functional sensitivity at milder stimulus intensities. We have previously implicated NompC, Trpm and Pkd2 in cold nociception, along with emerging evidence for proper chloride (Cl^−^) homeostasis being necessary for cold-evoked behaviors (Himmel, Sakurai, et al., 2021; Turner et al., 2016). It might be that at noxious cold temperature ranges that average spiking activity may be a result of TRP channels and Cl^−^ signaling, where GABA_B_-R2 or RyR function in patterning of cold evoked electrical activity in CIII md neurons. In our experimental paradigm, cold (10°C) stimulation of CIII md neurons consists of an initial rapid temperature decrease to target temperature and then steady state exposure for remaining of the recording. In control CIII md neurons, bursting response is observed during the initial rapid temperature decrease and then electrical activity transitions to primarily tonic spiking during steady state cold exposure. *RyR* depletion in CIII neurons leads to reduced bursting but increased tonic spiking, whereas RyR activation via ryanodine application facilitates bursting but has no effect on tonic spiking. However, for *GABA*_*B*_*-R2* knockdown in CIII md neurons, we observed a disorganized firing pattern, where the composition of bursting and tonic activity was altered compared to controls. Our analyses on bursting and tonic spike activity suggest an overall role of GABA_B_-R2 and RyR in establishing the normal cold-evoked firing pattern. Sensory perception results in a specifically patterned response that results in feature detections and behavioral choice selection (Allen & Marsat, 2018; Doron et al., 2014). For example, sensory coding in the rat barrel cortex is dependent on spike frequency, where irregular sensory stimuli led to greater behavioral response rate and high frequency microstimulations resulted in the most robust behavioral response (Doron et al., 2014). In weakly electric fish, *A. leptorhynchus*, electrosensory lateral line utilizes a dual strategy for stimulus discrimination, where heterogenous and graded changes in firing convey greater information content versus synchronous bursting which results in low information content regarding environmental cues relating to neighboring fish (Allen & Marsat, 2018). Additionally, in crickets, *T. oceanicus*, ultrasound sensitive AN2 interneurons have bursting spikes that are predictive of behavioral response (Marsat & Pollack, 2006). However, these investigations into neural encoding of sensory discrimination did not involve molecular bases of how action potential firing patterns at the level of individual neurons can encode stimulus evoked behavioral responses. Our findings present new possibilities for investigating the molecular bases of how sensory neuron spike patterning drives stimulus relevant behavioral responses.

A naïve hypothesis would be that loss of function in ER Ca^2+^ reuptake pathways may lead to increases in cytosolic Ca^2+^ levels thereby enhancing cold-evoked responses. However, knockdown of *SERCA, Stim*, or *Orai* in CIII md neurons led to significant reductions in cold-evoked CT responses. Dysregulated increases in cytosolic Ca^2+^ levels can lead to impaired activity of voltage gated Ca^2+^ channels, which are inactivated through Ca^2+^-calmodulin (CaM) complexes (Hering et al., 1997) and calcium imaging studies in *Drosophila* wing disc development revealed that loss of SERCA led to severely reduced Ca^2+^ dynamics (Brodskiy et al., 2019). Hyper-elevated Ca^2+^ can also negatively regulate TRP channel function, whereby Ca^2+^-CaM binding leads to channel inactivation (Gordon-Shaag et al., 2008), additionally, changes in Ca^2+^ gradients leading to localized outward Ca^2+^ current can cause long term inactivation of Pkd2 channels (DeCaen et al., 2016). Alternatively, excess Ca^2+^ might facilitate activation of Ca^2+^-activated K^+^ channels leading to disrupted firing patterns (Fakler & Adelman, 2008; Schumacher et al., 2001). How alterations in Ca^2+^ homeostasis lead to impaired cold insensitivity requires future investigations into interactions between Ca^2+^ signaling and PM localized ion channels. Alternatively, Ca^2+^ induced neuronal damage/degradation may also lead to impaired cold responses due to loss of function of ER Ca^2+^ reuptake machinery. High cytosolic Ca^2+^ could also lead to activation of Ca^2+^ sensitive proteases which have been shown to contribute to axonal degeneration (Vargas et al., 2015; Williams et al., 2014; Yang et al., 2013). Assessing roles of Ca^2+^ sensitive proteases, Calpains, which are ~10-fold upregulated in CIII md neurons compared to whole larva, in context of ER Ca^2+^ reuptake pathways and cold nociception may yield further insights into the importance of ER Ca^2+^ homeostasis. Overall, our data indicate that Ca^2+^ homeostasis is important for cold-evoked electrical patterning as well as behavioral output, where perturbations leading to reduced or excessive cytosolic Ca^2+^ result in disrupted cellular processes.

Excessive increases in cytosolic Ca^2+^ can be cytotoxic, thus Ca^2+^ homeostasis is tightly regulated through local organization of Ca^2+^ sensors, channels, and transporters. GPCRs, G-protein signaling machinery, CICR signaling machinery, and Ca^2+^ reuptake proteins are required for cold nociceptive behaviors in *Drosophila* larvae. Elevated Ca^2+^ levels can impact multiple downstream targets including Ca^2+^-activated K^+^ or Cl^−^ channels (Berg et al., 2012; Shah et al., 2006). Anoctamins are a highly conserved family of Ca^2+^ sensitive proteins that function as chloride ion channels, lipid scramblases, and/or molecular tethers for linking PM and ER membranes (Benarroch, 2017; Jha et al., 2019; Ye et al., 2018). Anoctamin 1, ANO1, is required for nociceptive heat sensing in DRGs and in nociceptive neurons. ANO1 is specifically activated by CICR induced cytosolic increase in Ca^2+^, which is dependent upon coupling with IP_3_R (Cho et al., 2012; Jin et al., 2013; Liu et al., 2010). We have recently implicated two *Drosophila* Anoctamin/TMEM16 channels, *subdued*, orthologous to human ANO1/2, and *white walker (wwk)*, orthologous to human ANO8, in *Drosophila* larval cold nociception (Himmel, Sakurai, et al., 2021). CIII md neurons lacking *subdued* or *wwk* have impaired cold-evoked electrical response, where knockdown of either gene led to reductions in tonic spiking activity which contrasts with our findings that *RyR* depleted neurons exhibit reduced bursting activity suggesting that cold-evoked electrical activity patterns (*i*.*e*. bursting vs. tonic spiking) can be parsed at the molecular level (Himmel, Sakurai, et al., 2021). Similar to findings in DRG cell cultures, CIII md neurons use excitatory Cl^−^ current physiology, where CIII-specific optogenetic activation of a light sensitive Cl^−^ channel, Aurora, evokes CT behavioral response (Cho et al., 2012; Himmel, Sakurai, et al., 2021). Additionally, increasing intracellular Cl^−^ concentration in CIII md neurons via *ncc69* overexpression, orthologous to human NKCC1, resulted in increased bursting activity. Cl^−^ ion homeostasis is not required for innocuous mechanical sensation but it is required for cold nociception, which suggests similar functional roles in sensory discrimination for GABA_B_-R2, IP_3_R and stimulus evoked Cl^−^ current in *Drosophila* multimodal CIII neurons (Himmel, Sakurai, et al., 2021). However, it remains unclear whether *Drosophila* Anoctamins function in conjunction with GPCRs, IP_3_R or RyR for stimulus evoked activation of cold sensitive pathways.

TRP channels are required for *Drosophila* larval cold nociceptive behaviors (Turner et al., 2016). Specifically, Pkd2 is required for cold-evoked increases in cytosolic Ca^2+^ and can confer cold sensitivity to non-cold sensitivity neurons. In mouse cell cultures, microdomains of Pkd2-IP_3_R interaction were required for CICR mechanism (Sammels et al., 2010). We explored whether a Pkd2 and IP_3_R genetically interact in the cold nociceptive pathway. Transheterozygous mutant analyses indicate that *Pkd2* and *IP*_*3*_*R* genetically interact and function synergistically in *Drosophila* larval cold nociception. Interestingly, Pkd2 function in *Drosophila* larva is not required for sensory discrimination between cold and innocuous touch (Turner et al., 2016). IP_3_R and Pkd2 synergistic function in cold nociception may be independent of GPCR mediated activation of IP_3_R pathway because at least our results suggest that GABA_B_ and IP_3_R are specifically required for cold sensing but not innocuous mechanical detection. Future investigations are required to clarify whether Pkd2, GABA_B_ and IP_3_R co-localize with one another to form microdomains and how these molecules collectively contribute to modality-specific sensory discrimination.

## 7 Author Contributions

Conceptualization: AAP and DNC. Methodology: AAP, AS, and DNC. Cold-plate assays: AAP; Innocuous touch assays: NJH. Optogenetics, AAP; Neural morphometric analyses: AAP. Calcium imaging: AAP. Electrophysiology: AS. Statistics and other formal analyses: AAP and AS. Writing – Original Draft: AAP and DNC; Writing – Review & Editing: AAP, AS, NJH, and DNC. Visualization: AAP and AS. Supervision: DNC. Funding acquisition: DNC.

## 9 Acknowledgments

We thank Romee Maitra for CaMPARI Sholl image processing. We thank Shatabdi Bhattacharjee and Dustin E.A. Moon in the Cox Lab for critical comments on the manuscript.

