## Supplemental Material for "Modality specific roles for metabotropic GABAergic signaling and calcium induced calcium release mechanisms in regulating cold nociception"

### *Supplementary Material*

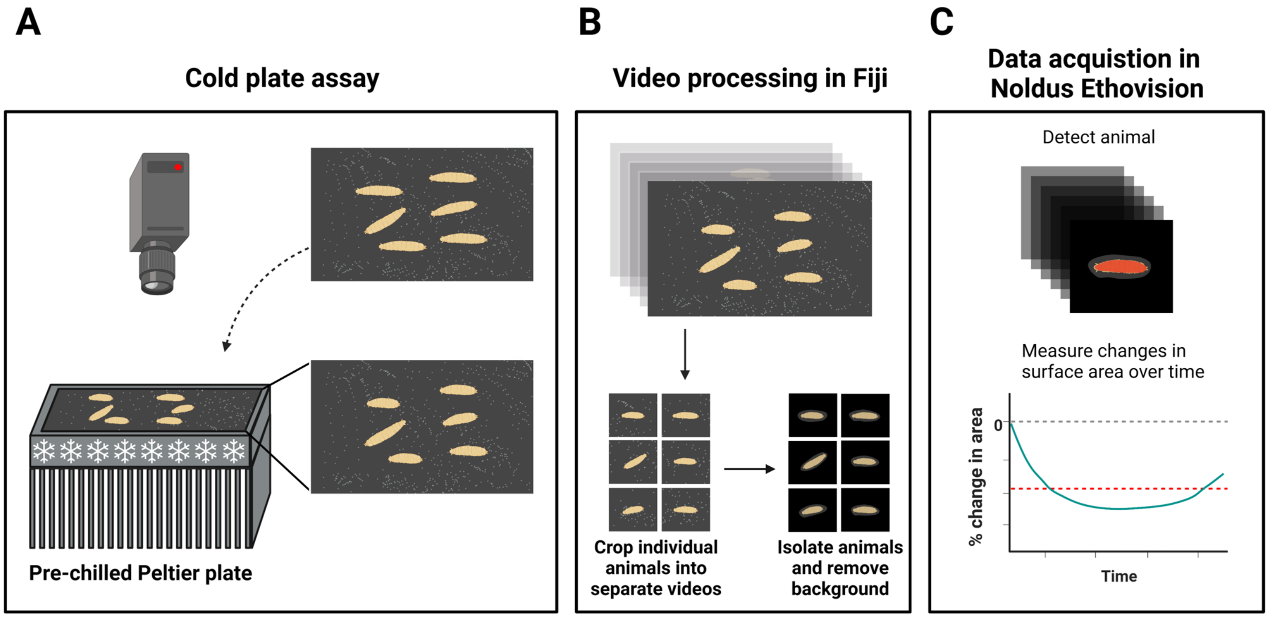


**Supplementary Figure 1: Cold plate assay schematic.** (**A**) Graphical representation of cold plate assay. Freely moving third instar larvae on black metal plate are exposed to noxious cold stimulus by placing the plate on pre-chilled Peltier surface. (**B**) Video processing pipeline in FIJI. Original video is trimmed from stimulus delivery to 5 seconds. Trimmed video is cropped into multiple videos to contain one larva per video. Background is removed from cropped videos for greater signal to noise ratio. (**C**) Acquisition of changes in larval surface area over time measurements are performed using Noldus Ethovision XT. Schematic created with BioRender.com.

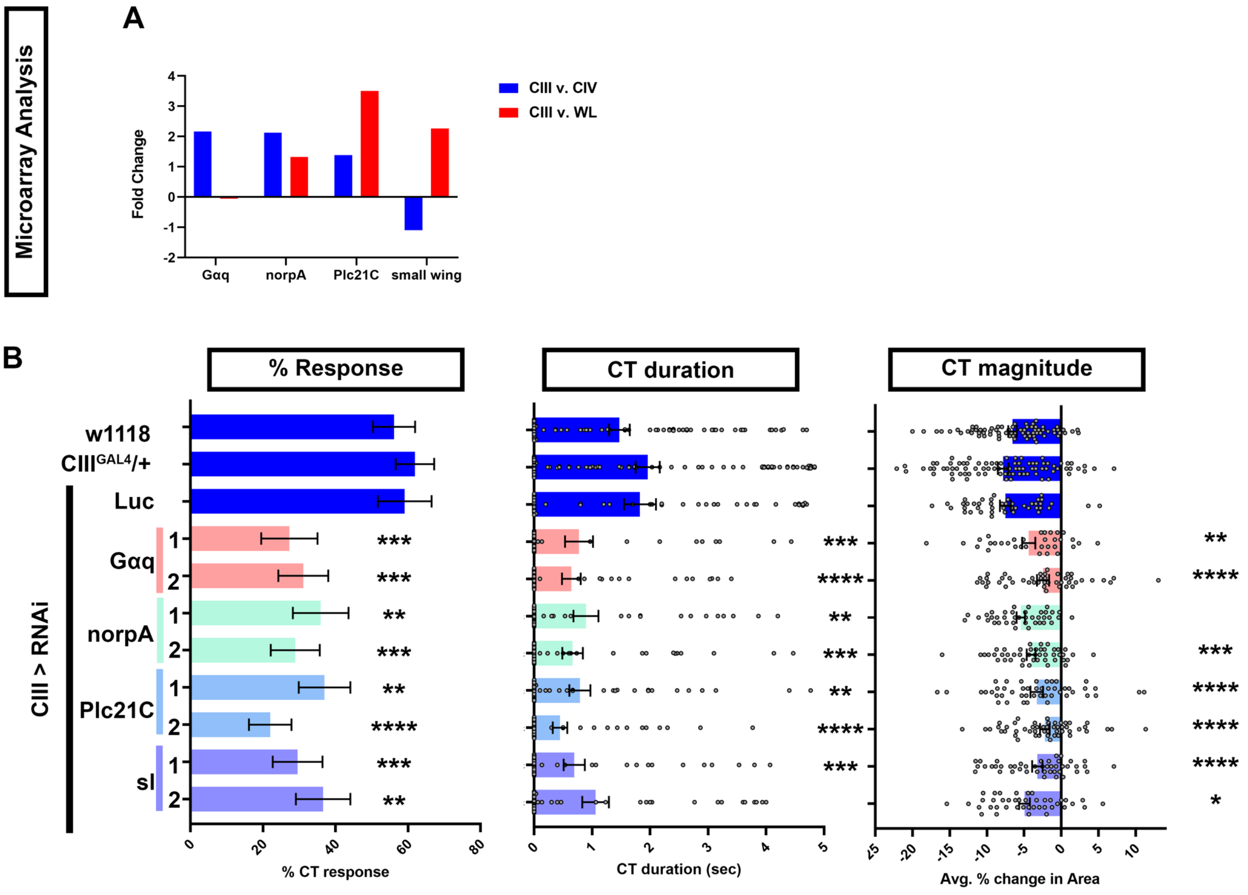


**Supplementary Figure 2: Gαq and phospholipase C required for cold evoked CT response in CIII md neurons**.

(**A**) Fold change in expression for $G_{\alpha q}$ and *PLCs* (*norpA*, *Plc21C*, and *small wing* (*sl*)) comparing CIII md neuron and CIV md neurons or whole larva (WL) microarray datasets. (**B**) Cold plate analysis of *Drosophila* larvae with CIII md neuron specific gene knockdown of $G_{\alpha q}$ and *PLCs*. We report %CT (left), CT duration in seconds (middle), CT magnitude (right). Control conditions include: *w1118*, *19-12*$^{GAL4}$/+, and *19-12>Luc*$^{RNAi}$. There is no statistical difference between the controls. Knockdown of $G_{\alpha q}$ and *PLCs* was tested with two independent RNAi lines and comparisons made to *19-12*$^{GAL4}$/+. $N_{Average}$ = 49. Significant differences indicated via asterisks, where *p<0.05, **p<0.01, ***p<0.001, ****p<0.0001.

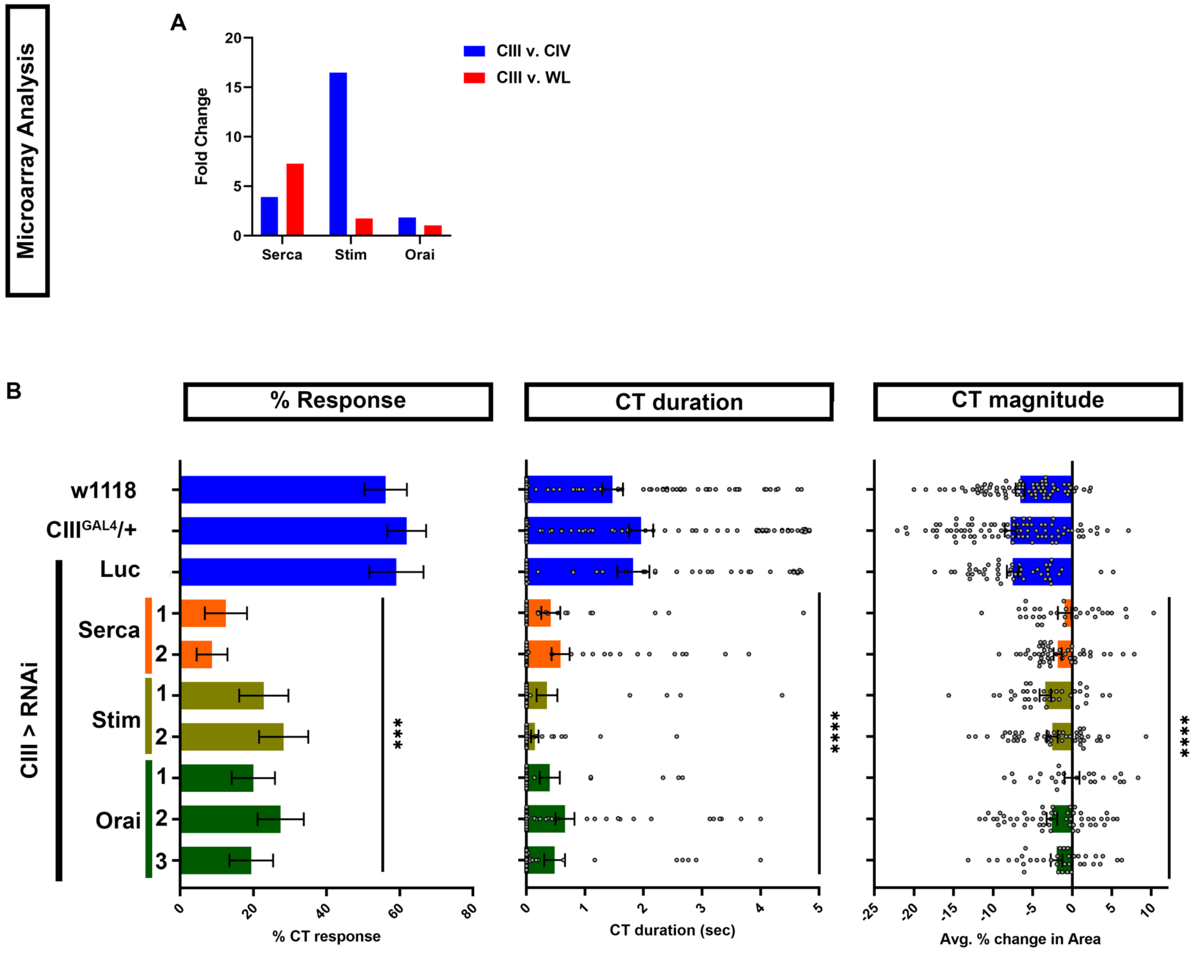


**Supplementary Figure 3: Requirement for Ca$^{2+}$ reuptake genes in cold evoked CT behavior.**

(**A**) Microarray analysis of *SERCA*, *Stim* and *Orai* in CIII md neurons compared to CIV md neurons or whole larva (WL). (**B**) Ca$^{2+}$ reuptake gene knockdown specifically in CIII md neurons using *19-12*$^{GAL4}$ driver. We report %CT (left), CT duration in seconds (middle), CT magnitude (right). Control conditions include: *w1118*, *19-12*$^{GAL4}$*/+,* and *19-12>Luc*$^{RNAi}$. There is no statistical difference between the controls. Knockdown of *SERCA*, *Stim* and *Orai* was tested with at least two independent RNAi lines and comparisons made to *19-12*$^{GAL4}$/+. N$_{Average}$ = 47. Significant differences indicated via asterisks, where *p<0.05, **p<0.01, ***p<0.001, ****p<0.0001.

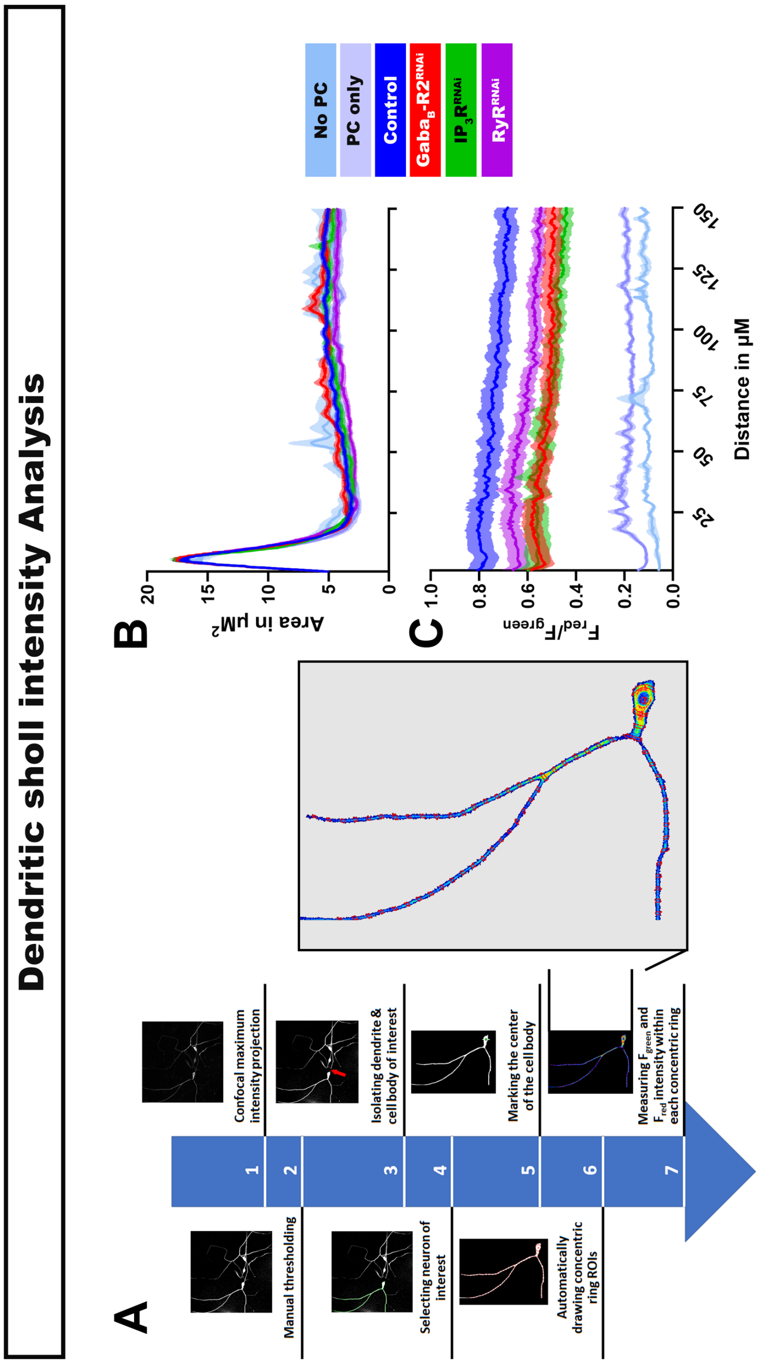


**Supplementary Figure 4: 2D Sholl intensity analysis of dendritic CIII md neuronal CaMPARI2 response.**

(**A**) Schematic of custom FIJI macro script for rapidly removing background, selecting cell body and dendrites of interest, automatic generation of sholl ROIs, and measuring area normalized $F_{red}$ and $F_{green}$ intensities. (**B**) Average area in µm$^2$ for individual ROIs as function of distance from the center of the cell body. (**C**) Area normalized average $F_{red}/F_{green}$ ratio of CaMPARI2 plotted as a function of distance away from the cell body. $N_{Average} = 30$.

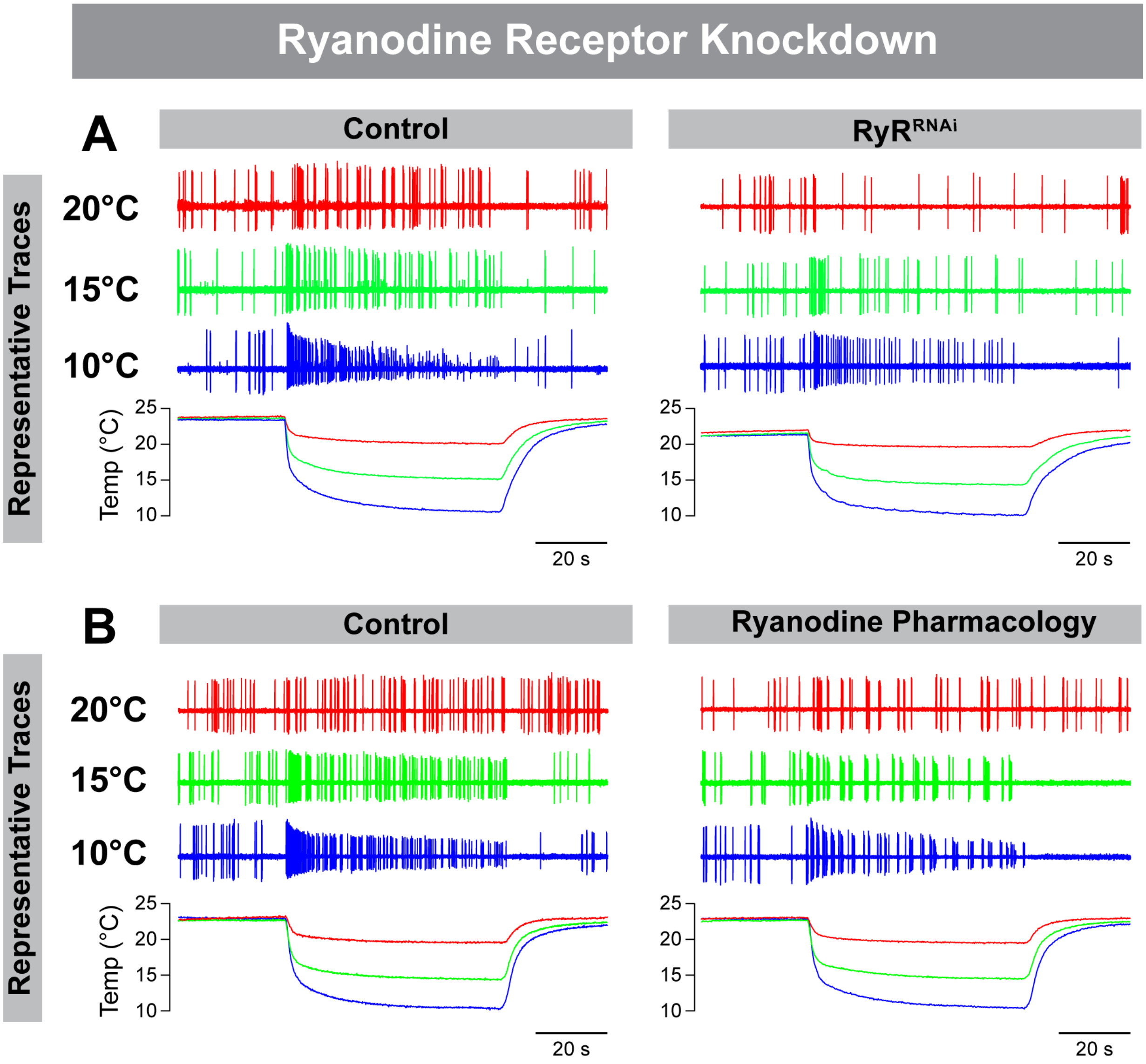


**Supplementary Figure 5: Representative extracellular recording traces CIII md neurons.**

Representative physiological firing traces for each stimulus temperature (20°C, 15°C, 10°C) and below shows real time temperature exposure regimes of *Drosophila* larval filet. (**A**) *RyR* knockdown compared to controls. (**B**) Pharmacological application of ryanodine ($10^{-6}$ M) compared to controls.

**Table 1: List of Drosophila melanogaster strains**

| Designation | Source or reference | Identifier |
|---|---|---|
| ***w1118*** | BDSC | 3605 |
| ***19-12$^{GAL4}$*** | Yan et al. (2012). *Nature* 493:221-225 | |
| ***GMR83B04$^{GAL4}$*** | BDSC | 41309 |
| ***nompC$^{GAL4}$*** | BDSC | 36361 |
| ***UAS-CD4-tdGFP*** | BDSC | 35836 |
| ***UAS-CaMPARI2*** | BDSC | 78317 |
| ***UAS-IVS-GCaMP6m*** | BDSC | 42748 |
| ***UAS-ChETA::YFP*** | BDSC | 36495 |
| ***Luc RNAi*** | BDSC | 31603 |
| ***GABA-B-R1 RNAi - 1*** | BDSC | 28353 |
| ***GABA-B-R1 RNAi - 2*** | BDSC | 51817 |
| ***GABA-B-R2 RNAi - 1*** | BDSC | 50608 |
| ***GABA-B-R2 RNAi - 2*** | BDSC | 27699 |
| ***GABA-B-R3 RNAi - 1*** | BDSC | 26729 |
| ***GABA-B-R3 RNAi - 2*** | BDSC | 50622 |
| ***GABA-B-R1 MiMIC*** | BDSC | 44860 |
| ***GABA-B-R1 KO - CRISPR*** | BDSC | 84503 |
| ***GABA-B-R1 KI - CRISPR*** | BDSC | 84701 |
| ***GABA-B-R2 MiMIC*** | BDSC | 59503 |
| ***GABA-B-R2 KO - CRISPR*** | BDSC | 84504 |
| ***GABA-B-R2 KI - CRISPR*** | BDSC | 84634 |
| ***GABA-B-R3 MiMIC*** | BDSC | 23554 |
| ***GABA-B-R3 KO - CRISPR*** | BDSC | 84505 |
| ***GABA-B-R3 KI - CRISPR*** | BDSC | 84635 |
| ***Gαq RNAi - 1*** | BDSC | 36820 |
| ***Gαq RNAi - 2*** | BDSC | 36775 |
| ***norpA RNAi - 1*** | BDSC | 31113 |
| ***norpA RNAi - 2*** | BDSC | 31197 |
| ***Plc21C RNAi - 1*** | BDSC | 31269 |
| ***Plc21C RNAi - 2*** | BDSC | 33719 |
| ***sl RNAi - 1*** | BDSC | 32385 |
| ***sl RNAi - 2*** | BDSC | 32906 |
| ***IP3R RNAi - 1*** | BDSC | 51795 |
| ***IP3R RNAi - 2*** | BDSC | 51686 |
| ***IP3R RNAi - 3*** | BDSC | 25927 |
| ***RyR RNAi - 1*** | BDSC | 31695 |

| | | |
|---|---|---|
| ***RyR RNAi - 2*** | BDSC | 31540 |
| ***RyR RNAi - 3*** | BDSC | 28919 |
| ***RyR RNAi - 4*** | BDSC | 29445 |
| ***IP3R 90B0*** | BDSC | 30737 |
| ***IP3R ug3*** | BDSC | 30738 |
| ***IP3r ka901*** | BDSC | 30741 |
| ***RyR 16*** | BDSC | 6812 |
| ***RyR Q3878X*** | BDSC | 5501 |
| ***RyR Y4452X*** | BDSC | 5507 |
| ***Serca RNAi - 1*** | BDSC | 25928 |
| ***Serca RNAi - 2*** | BDSC | 44581 |
| ***Orai RNAi - 1*** | BDSC | 53333 |
| ***Orai RNAi - 2*** | VDRC | GD12221 |
| ***Stim RNAi - 1*** | BDSC | 27263 |
| ***Stim RNAi - 2*** | BDSC | 51685 |
| ***Stim RNAi - 3*** | BDSC | 52911 |
| ***PKD2[1] mutant*** | BDSC | 24495 |
